# Predicting bioprocess targets of chemical compounds through integration of chemical-genetic and genetic interaction networks

**DOI:** 10.1101/111252

**Authors:** Scott W. Simpkins, Justin Nelson, Raamesh Deshpande, Sheena C. Li, Jeff S. Piotrowski, Erin H. Wilson, Abraham A. Gebre, Reika Okamoto, Mami Yoshimura, Michael Costanzo, Yoko Yashiroda, Yoshikazu Ohya, Hiroyuki Osada, Minoru Yoshida, Charles Boone, Chad L. Myers

## Abstract

Chemical-genetic interactions – observed when the treatment of mutant cells with chemical compounds reveals unexpected phenotypes – contain rich functional information linking compounds to their cellular modes of action. To systematically identify these interactions, an array of mutants is challenged with a compound and monitored for fitness defects, generating a chemical-genetic interaction profile that provides a quantitative, unbiased description of the cellular function(s) perturbed by the compound. Genetic interactions, obtained from genome-wide double-mutant screens, provide a key for interpreting the functional information contained in chemical-genetic interaction profiles. Despite the utility of this approach, integrative analyses of genetic and chemical-genetic interaction networks have not been systematically evaluated. We developed a method, called CG-TARGET (Chemical Genetic Translation via A Reference Genetic nETwork), that integrates large-scale chemical-genetic interaction screening data with a genetic interaction network to predict the biological processes perturbed by compounds. CG-TARGET compared favorably to a baseline enrichment approach across a variety of benchmarks, achieving similar accuracy while substantially improving the ability to control the false discovery rate of biological process predictions. We applied CG-TARGET to a recent screen of nearly 14,000 chemical compounds in *Saccharomyces cerevisiae*, integrating this dataset with the global *S. cerevisiae* genetic interaction network to prioritize over 1500 compounds with high-confidence biological process predictions for further study. Upon investigation of the compatibility of chemical-genetic and genetic interaction profiles, we observed that one-third of observed chemical-genetic interactions contributed to the highest-confidence biological process predictions and that negative chemical-genetic interactions overwhelmingly formed the basis of these predictions. We present here a detailed characterization of the CG-TARGET method along with experimental validation of predicted biological process targets, focusing on inhibitors of tubulin polymerization and cell cycle progression. Our approach successfully demonstrates the use of genetic interaction networks in the functional annotation of compounds to biological processes.

## Author Summary

Understanding how chemical compounds affect biological systems is of paramount importance as pharmaceutical companies strive to develop life-saving medicines, governments seek to regulate the safety of consumer products and agrichemicals, and basic scientists continue to study the fundamental inner workings of biological organisms. One powerful approach to characterize the effects of chemical compounds in living cells is chemical-genetic interaction screening. Using this approach, a collection of cells – each with a different defined genetic perturbation – is tested for sensitivity or resistance to the presence of a compound, resulting in a quantitative profile describing the functional effects of that compound on the cells. The work presented here describes our efforts to integrate compounds’ chemical-genetic interaction profiles with reference genetic interaction profiles containing information on gene function to predict the cellular processes perturbed by the compounds. We focused on specifically developing a method that could scale to perform these functional predictions for large collections of thousands of screened compounds and robustly control the false discovery rate. With chemical-genetic and genetic interaction screens now underway in multiple species including human cells, the method described here can be generally applied to enable the characterization of compounds’ effects across the tree of life.

## Introduction

The ability to discover chemical compounds with desirable and interesting biological activity is essential for understanding how compounds and biological systems interact. Chemical-genetic interaction screening provides a means to characterize the biological activity of compounds in an unbiased manner by measuring the response of defined gene mutants to these molecules [1–8]. A chemical-genetic interaction profile refers to the set of gene mutations that confer sensitivity (a negative chemical-genetic interaction) or resistance (a positive interaction) to a compound and provides functional insights into the compound’s mode(s) of action.

Similar to chemical-genetic interactions, genetic interactions identify pairs of gene mutations whose combined effects are more or less severe than expected given the phenotypes of the individual mutants. In *S. cerevisiae*, the vast majority of all possible gene double-mutant pairs have been constructed and scored for fitness-based genetic interactions, yielding a global compendium of genome-wide genetic interaction profiles that quantitatively describe each gene’s function. Similarity between two genes’ genetic interaction profiles implies that these genes perform similar cellular functions, enabling the functional annotation of previously unannotated genes and the construction of a global hierarchy of cellular function [5,9].

Chemical-genetic and genetic interaction profiles derived from fitness measurements contain analogous functional information on the cellular effects of chemicals and gene mutations, respectively. Similarity between these two types of profiles therefore implies that the respective chemical(s) and gene mutation(s) perturb similar functions in the cell, which means that a compound’s chemical-genetic interaction profile should resemble the genetic interaction profile(s) of its cellular target or target processes (Fig 1) [2,5]. The global genetic interaction network in *S. cerevisiae* therefore provides a resource for interpreting chemical-genetic interaction profiles across a broad range of cellular function. Importantly, this approach to interpretation does not depend on reference chemical-genetic interaction profiles and thus enables the discovery of compounds with novel modes of action.

**Figure 1.**
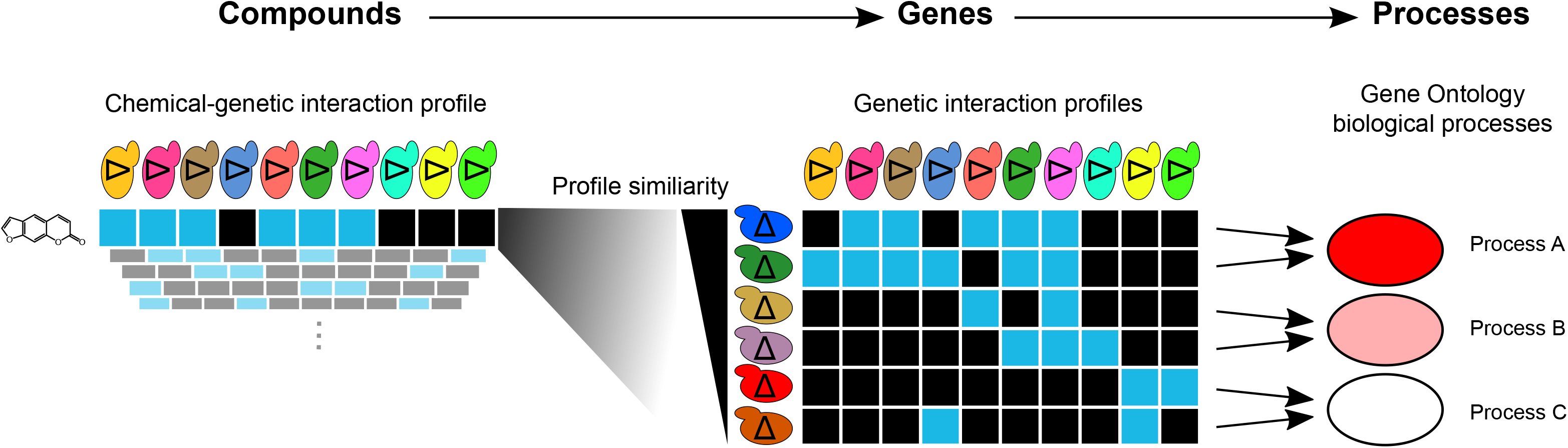
Overview of the integration of chemical-genetic and genetic interaction networks for bioprocess target prediction using CG-TARGET. Chemical-genetic interaction profiles, obtained by measuring the sensitivity or resistance of a library of gene mutants to a chemical compound, are compared against genetic interaction profiles consisting of double mutant interaction scores. The resulting similarities are aggregated at the level of biological processes to predict the bioprocess(es) perturbed by the compound. Better agreement between chemical-genetic and genetic interaction profiles leads to stronger bioprocess predictions. Each blue box represents a negative chemical-genetic (i.e. sensitivity) or genetic interaction, while each black box represents the absence of an interaction. Stronger bioprocess predictions are depicted with a darker red.

Recent advances in DNA sequencing technology have paved the way for high-throughput chemical-genetic interaction screening via multiplexed analysis of pooled, genetically-barcoded mutant libraries grown in the presence of compound [6, 7, 10]. This would enable, for example, functional profiling of compounds earlier in the drug discovery process, with insights from these screens providing a means to prioritize compounds before investing resources into their development as drugs. Despite the recent generation of thousands of chemical-genetic interaction profiles across multiple studies [6,7] and the profound opportunities for genetic interaction-powered functional characterization of thousands of novel compounds, the integration of chemical-genetic and genetic interaction profiles has only been performed in the context of relatively small studies [2,5]. A systematic investigation using a large-scale chemical-genetic interaction dataset is therefore necessary to assess the compatibility between chemical-genetic and genetic interaction profiles, with an emphasis on the ability of a genetic interaction-based method to control the false discovery rate (of critical importance in high-throughput chemical screening) and thereby prioritize compounds with the highest-confidence predictions.

Here, we present the use of genetic interaction profiles to systematically interpret chemical-genetic interaction profiles on a large scale. Specifically, we developed a computational method, called CG-TARGET (Chemical Genetic Translation via A Reference Genetic nETwork), that integrates chemical-genetic and genetic interaction profiles to predict the biological processes perturbed by compounds. We applied this method to a high-throughput chemical-genetic interaction screen of nearly 14,000 compounds in *S. cerevisiae* [11], using profiles from the global yeast genetic interaction network [5,9] to interpret the chemical-genetic interaction profiles. CG-TARGET recapitulated known information for well-characterized compounds and showed a marked improvement in the ability to control the false discovery rate for novel compound mode-of-action discovery compared to a baseline approach. Additionally, we experimentally validated two different mode-of-action predictions, one in an *in vitro* system using mammalian proteins, confirming both the accuracy of the predictions for novel compounds and the potential to translate these predictions across species. CG-TARGET is available, free for academic use, at https://github.com/csbio/CG-TARGET.

## Results

### Overview of datasets used in this study

We obtained chemical-genetic interaction profiles from a recent large-scale chemical-genetic interaction screen in *S. cerevisiae* [11]. This screen consisted of two batches, the first of which containing 9850 compounds from the RIKEN Natural Product Depository [12] (the “RIKEN” screen) and the second containing 4116 compounds from the NCI Open Chemical Repository’s compound libraries, the NIH Clinical Collection, and GlaxoSmithKline’s Published Kinase Inhibitor Set (the “NCI/NIH/GSK” screen) [13]. The compounds in the RIKEN screen consisted primarily of natural products and natural product derivatives – most of which were previously uncharacterized – and ~200 approved drugs and chemical probes, a subset of which we used to assess the performance of CG-TARGET as their modes of action in yeast are well-characterized. The compounds in the NCI/NIH/GSK screen were more studied – having been tested against the NCI-60 cancer cell line panel (the NCI collections), tested in clinical trials (the NIH Clinical Collection) or designed and characterized as inhibitors against human kinases (GSK) – but many of these compounds’ specific modes of action remain uncharacterized. The final datasets consisted of 8418 chemical-genetic interaction profiles from the RIKEN screen and 3565 from the NCI/NIH/GSK screen, which were obtained using a diagnostic set of approximately 300 haploid gene deletion mutants that were optimally selected to capture most of the information in the complete *S. cerevisiae* non-essential deletion collection [11,14]. Both datasets also contained a large set of experimental control profiles (5724 and 2128 for the RIKEN and NCI/NIH/GSK screens, respectively), in which the yeast were only treated with the solvent control (DMSO). Each profile contains z-scores that reflect the deviation of each strain’s observed fitness from expected fitness in the presence of a compound.

Genetic interaction profiles were obtained from a recently assembled, genome-wide compendium of genetic interaction profiles in *S. cerevisiae* [5]. These profiles were generated by systematically constructing and analyzing the fitness of haploid double mutant strains and consist of epsilon scores that reflect the deviation of each double mutant’s observed fitness from that expected given the single mutant fitness values, assuming a multiplicative null model [15]. The construction of each profile involved crossing the mutant for the “query” gene into a genome-wide array of mutants, and we mapped the query genes to Gene Ontology biological process terms [16,17] to define the bioprocess targets of compounds. Profiles were filtered to the ~35% with the highest signal (see Materials and Methods).

### Predicting perturbed bioprocesses from chemical-genetic interaction profiles

We developed CG-TARGET (Chemical Genetic Translation via A Reference Genetic nETwork) to predict the biological processes perturbed by compounds in our recently-generated dataset of ~12,000 chemical-genetic interaction profiles (Fig 1). CG-TARGET requires three input datasets: 1) chemical-genetic interaction profiles; 2) genetic interaction profiles; and 3) a mapping from the query genes in the genetic interaction profiles to gene sets representing coherent bioprocesses. Predicting the bioprocesses perturbed by a particular compound involves four distinct steps. First, a control set of resampled chemical-genetic interaction profiles is generated, each of which consists of one randomly-sampled interaction score per gene mutant across all compound treatment profiles in the chemical-genetic interaction dataset; these profiles thus provide a means to account for variance in each mutant strain observed upon treatment with bioactive compound but not upon treatment with experimental controls (DMSO with no active compound). Second, scores reflecting both the strength of each compound’s chemical-genetic interaction profile and its similarity to the profile of each gene mutant are obtained by computing an inner product between all chemical-genetic interaction profiles (comprising compound treatment, experimental control, and random profiles) and all *L*_2_-normalized query genetic interaction profiles. Third, these “gene-level” prediction scores are aggregated into bioprocess predictions; a z-score and empirical p-value for each compound-bioprocess prediction are obtained by mapping the gene-level prediction scores to the genes in the bioprocess of interest and comparing these scores to those from shuffled gene-level prediction scores and to distributions of the scores derived from experimental control and resampled profiles. Finally, the false discovery rates for these predictions are estimated by comparing, across a range of significance thresholds, the frequency at which experimental control and randomly resampled profiles predict bioprocesses versus that of compound treatment profiles (see Materials and Methods).

### Application to and evaluation on large-scale chemical-genetic interaction data

To provide a baseline method for benchmarking the performance of CG-TARGET on these large screens, we implemented a simple, enrichment-based approach for predicting bioprocess-level targets. The enrichment-based approach was designed to predict bioprocess-level targets by testing for the enrichment of GO biological processes among the top-*n* gene-level prediction scores for each compound. For the following comparisons, CG-TARGET was compared to top-20 enrichment, which showed the best overall performance across a range of values of *n* (Fig S1).

We applied CG-TARGET to the RIKEN and NCI/NIH/GSK chemical-genetic interaction screens, identifying 848 out of 8418 compounds (10%) from the RIKEN screen and 705 of 3565 compounds (20%) from the NCI/NIH/GSK screen with at least one prediction that achieved false discovery rates of 25 and 27%, respectively (referred to as “high-confidence” compounds and predictions) (Table 1, Fig 2). In all cases, the false discovery rates derived from resampled profiles were more conservative than those derived from experimental controls, suggesting that some sources of variance in each gene mutant’s interaction scores arose only upon treatment with compound and therefore could not be corrected using only solvent controls. Focusing on the results from the RIKEN screen, CG-TARGET substantially outperformed the baseline method with regard to the number of compounds that possessed at least one high-confidence bioprocess prediction (FDR ≤ 25%). Compared to the 848 high-confidence compounds identified by CG-TARGET, top-20 enrichment only identified seven compounds that met this confidence threshold, and zero with a false discovery rate less than 21% (Fig 3A).

**Table 1.**
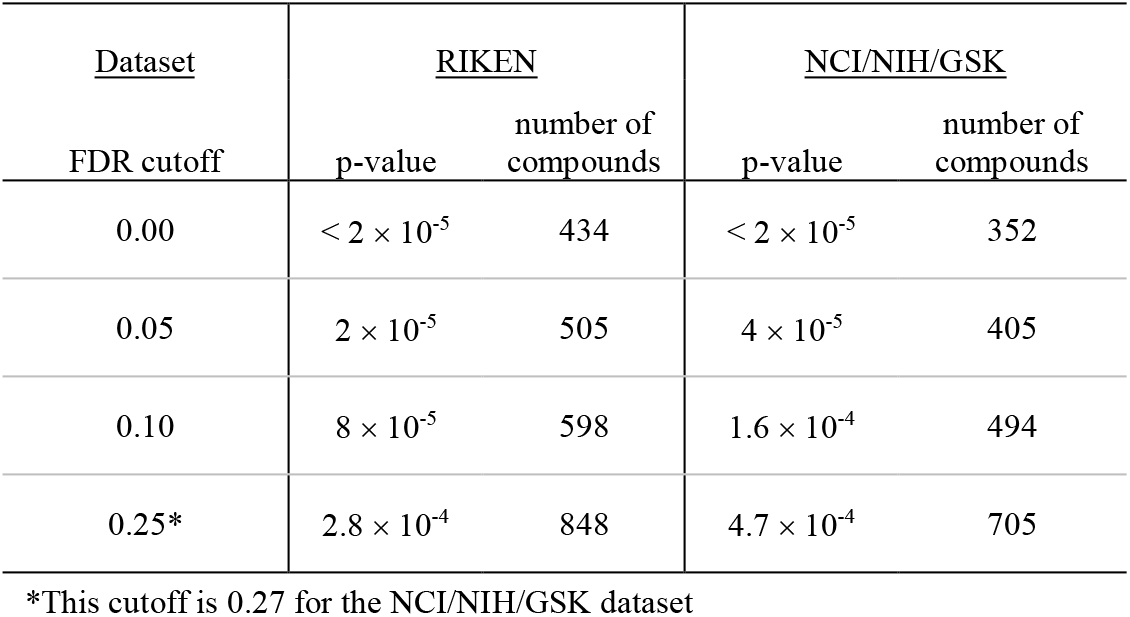
The number of compounds discovered at selected false discovery rates upon application of CG-TARGET to data from two large-scale chemical-genetic interaction screens. The “RIKEN” screen consisted of 8418 total compounds from the RIKEN Natural Product Depository, and the “NCI/NIH/GSK” consisted of 3565 compounds across 6 chemical compound collections from the National Cancer Institute, National Institutes of Health, and GlaxoSmithKline.

**Figure 2.**
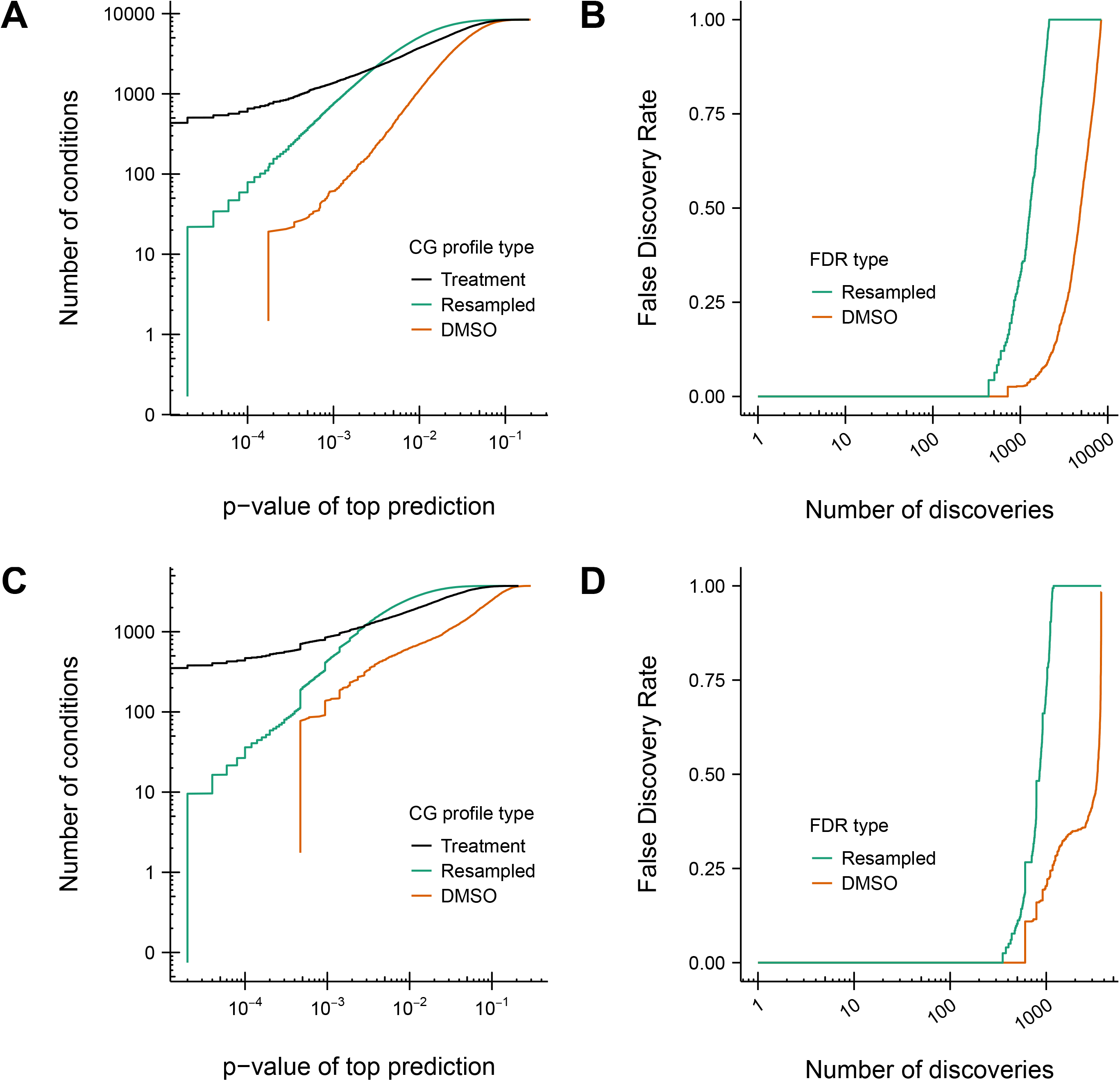
Rate of compound discovery and control of the false discovery rate for the prediction of bioprocesses from chemical-genetic interaction profiles. Perturbed bioprocesses were predicted using CG-TARGET for compounds, negative controls (DMSO), and resampled chemical-genetic interaction profiles from the RIKEN and NCI/NIH/GSK datasets. (A) The number of compounds, experimental controls, and randomly resampled chemical-genetic interaction profiles discovered with at least one bioprocess prediction passing the given significance thresholds, for the RIKEN dataset. (B) DMSO and resampled profile-derived estimates of the false discovery rate of biological process predictions, for the RIKEN dataset, given the number of discovered compounds. Values were calculated from (A). (C-D) Same as (A-B), respectively, but for the NCI/NIH/GSK dataset.

**Figure 3.**
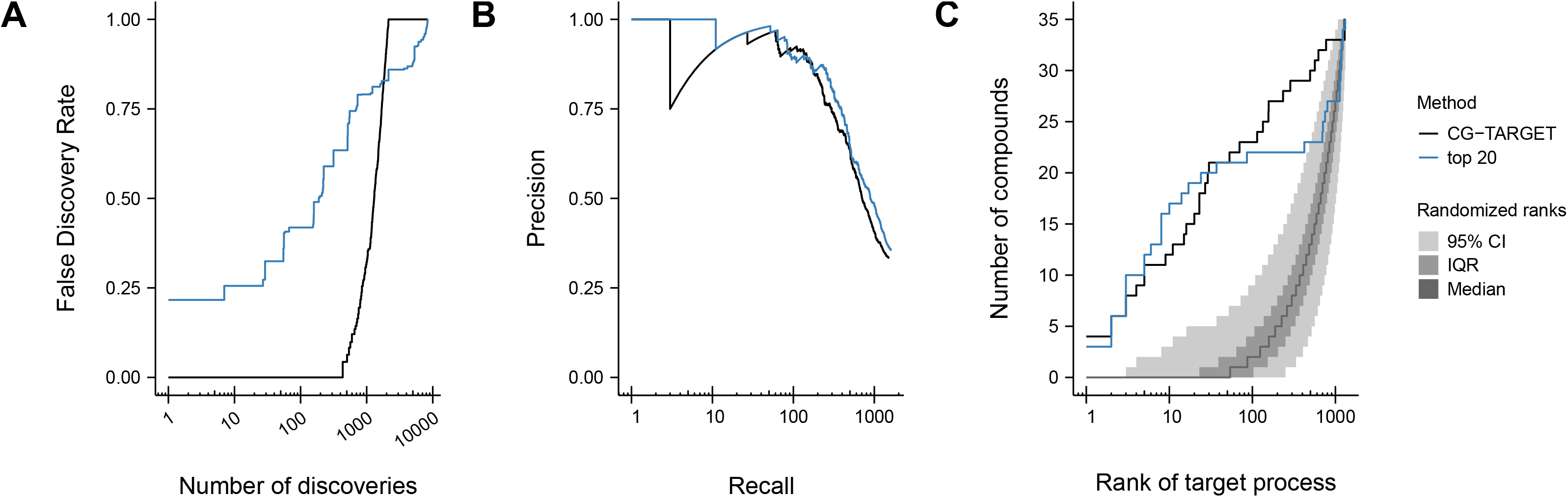
Performance comparison of CG-TARGET versus a baseline enrichment approach. Perturbed bioprocesses were predicted using both CG-TARGET and a method that calculated enrichment on the set of each compound’s 20 most similar genetic interaction profiles (“top 20”). (A) Bioprocess prediction false discovery rate estimates derived from resampled chemical-genetic interaction profiles, performed on compounds from the RIKEN dataset. (B) Precision-recall analysis of the ability to recapitulate gold-standard annotations within the set of top bioprocess predictions for ~4500 simulated compounds. Each simulated compound was designed to target one query gene in the genetic interaction network and thus inherited goldstandard biological process annotations from its target gene. (C) For each of 35 well-characterized compounds in the RIKEN dataset with literature-derived, gold-standard biological process annotations, we determined the rank of its gold-standard bioprocess within its list of predictions. The number of compounds for which a given rank (or better) was achieved is plotted. The grey ribbons represent the median, interquartile range (25^th^ to 75^th^ percentiles), and 95% confidence interval of 10,000 rank permutations.

CG-TARGET was also benchmarked against the baseline method using two different measures of prediction accuracy. The first accuracy-based evaluation was performed on genetic interaction profiles with added noise, which provided a means to both simulate chemical-genetic interaction profiles and annotate them with gold-standard GO biological process annotations for evaluation. For the second accuracy-based evaluation, we curated a set of gold-standard compound-bioprocess annotations from the literature for 35 compounds from the RIKEN screen and evaluated the ranks of the gold-standard bioprocesses within each compound’s list of bioprocess predictions.

CG-TARGET performed comparably to the best-performing enrichment-based methods using our measures of accuracy. This is first shown in the evaluation of these methods’ respective abilities to predict a gold-standard annotated bioprocess as the top prediction for each simulated chemical-genetic interaction profile. Specifically, CG-TARGET performed nearly as well as the top-20 enrichment-based method across both low and high recall values (Fig 3B). Both methods captured a gold-standard annotation as the top predicted bioprocess for approximately 34% of the simulated compounds (33.4% and 35.6% for CG-TARGET and top-20 enrichment, respectively), which represented more than a 22-fold enrichment over the background expectation of 1.5% (the average number of gold-standard bioprocess annotations per simulated compound divided by the number of bioprocesses).

Secondly, for the 35 gold-standard compounds with known target bioprocesses, we observed that both methods captured the gold-standard bioprocess for 6 and 21 (out of 35) compounds above ranks of 2 and 40 (out of 1329), respectively, with slightly decreased performance for CG-TARGET between these rank thresholds (Fig 3C, Table 2). The significance of these rank values was evaluated by randomizing the order of each compound’s bioprocess predictions 10,000 times and recalculating the ranks. Both methods achieved similar results in this respect, with CG-TARGET and the top-20 enrichment method respectively identifying 22 and 21 gold-standard compounds with significantly better ranks than the random expectation. CG-TARGET and top-20 enrichment also performed similarly when comparing the “effective rank” of each compound’s gold-standard bioprocess, with CG-TARGET and top-20 enrichment respectively identifying 20 and 22 compounds for which the gold-standard or a closely-related bioprocess achieved a rank of 5 or better.

**Table 2.** Evaluation of predictions made by CG-TARGET, and comparison to a baseline enrichment approach, for literature-derived, gold-standard compound-process annotations. The target bioprocess rank was determined by its position in the list of all bioprocess predictions for each gold-standard compound, with the significance computed empirically by shuffling the bioprocesses and re-computing the rank (bold p-values indicate significance, p < 0.05). Asterisks indicate cases in which the false discovery rate of the goldstandard compound-process prediction was less than 25%. The “top-20 enrichment” approach was selected as a baseline for comparison. The “effective rank” of a compound-bioprocess prediction represents the top rank within the compound’s list of predictions among bioprocesses that are similar to the original bioprocess.

| Compound | GO ID | GO term | CG-TARGET |  |  | top-20 enrichment |  |  |
| --- | --- | --- | --- | --- | --- | --- | --- | --- |
|  |  |  | Target process rank | Rank significance | Effective rank | Target process rank | Rank significance | Effective rank |
| 5-Fluorocytosine | GO:0032774 | RNA biosynthetic process | 27 | **0.0208** | 2 | 3 | **0.0027** | 1 |
| Aclacinomycin A | GO:0071103 | DNA conformation change | 1 | **\*0.0009** | 1 | 86 | 0.0643 | 2 |
| Acriflavine | GO:0006259 | DNA metabolic process | 30 | **\*0.0238** | 1 | 5 | **0.0042** | 1 |
| Benomyl | GO:0007017 | microtubule-based process | 2 | **\*0.0015** | 2 | 8 | **0.0056** | 2 |
| Blasticidin S | GO:0006412 | translation | 772 | 0.5842 | 57 | 1311 | 0.9883 | 247 |
| Bortezomib | GO:0030163 | protein catabolic process | 3 | **0.0026** | 1 | 8 | **0.0084** | 1 |
| Brefeldin A | GO:0006888 | ER to Golgi vesicle-mediated transport | 565 | 0.4207 | 32 | 1172 | 0.8818 | 169 |
| Caffeine | GO:0031929 | TOR signaling cascade | 1 | **\*0.0007** | 1 | 1 | **0.0007** | 1 |
| Calcofluor White | GO:0071554 | cell wall organization or biogenesis | 624 | 0.4675 | 90 | 1127 | 0.8526 | 176 |
| Camptothecin | GO:0071103 | DNA conformation change | 16 | **\*0.0114** | 4 | 6 | **0.0040** | 1 |
| Cisplatin | GO:0006260 | DNA replication | 134 | 0.1018 | 23 | 10 | **0.0071** | 1 |
| Daunorubicin | GO:0006260 | DNA replication | 70 | 0.0530 | 21 | 1210 | 0.9092 | 178 |
| FK228 | GO:0006325 | chromatin organization | 23 | **\*0.0169** | 2 | 17 | **0.0131** | 2 |
| Fluconazole | GO:0008202 | steroid metabolic process | 114 | 0.0870 | 12 | 708 | 0.5333 | 187 |
| Furazolidone | GO:0006260 | DNA replication | 20 | **\*0.0148** | 4 | 5 | **0.0034** | 1 |
| Gramicidin S | GO:0071554 | cell wall organization or biogenesis | 286 | 0.2186 | 39 | 1151 | 0.8705 | 173 |
| Griseofulvin | GO:0007017 | microtubule-based process | 1291 | 0.9718 | 227 | 750 | 0.5673 | 216 |
| Haloperidol | GO:0008202 | steroid metabolic process | 5 | **\*0.0035** | 2 | 37 | **0.0279** | 6 |
| Hedamycin | GO:0006281 | DNA repair | 4 | **\*0.0029** | 1 | 3 | **0.0022** | 1 |
| Hydroxyurea | GO:0006260 | DNA replication | 29 | **0.0239** | 6 | 1236 | 0.9269 | 1 |
| Itraconazole | GO:0008202 | steroid metabolic process | 234 | 0.1786 | 29 | 696 | 0.5239 | 193 |
| Latrunculin B | GO:0007010 | cytoskeleton organization | 11 | **\*0.0083** | 1 | 8 | **0.0068** | 2 |
| Micafungin | GO:0071554 | cell wall organization or biogenesis | 495 | 0.3718 | 47 | 1134 | 0.8577 | 150 |
| Mitomycin | GO:0006260 | DNA replication | 15 | **0.0104** | 4 | 2 | **0.0014** | 1 |
| MMS | GO:0006281 | DNA repair | 3 | **\*0.0022** | 1 | 3 | **0.0022** | 1 |
| Mycophenolic acid | GO:0006259 | DNA metabolic process | 1 | **\*0.0006** | 1 | 3 | **0.0025** | 1 |
| Nigericin | GO:0048193 | Golgi vesicle transport | 157 | 0.1158 | 13 | 1 | **0.0007** | 1 |
| Nocodazole | GO:0007017 | microtubule-based process | 2 | **\*0.0015** | 2 | 14 | **0.0100** | 3 |
| Oligomycin A | GO:0009268 | response to pH | 9 | **0.0075** | 2 | 2 | **0.0012** | 1 |
| Podophyllotoxin | GO:0007017 | microtubule-based process | 53 | **0.0411** | 6 | 800 | 0.6038 | 157 |
| Polyoxin D | GO:0071554 | cell wall organization or biogenesis | 1302 | 0.9788 | 225 | 1168 | 0.8828 | 173 |
| Rapamycin | GO:0031929 | TOR signaling cascade | 156 | 0.1140 | 8 | 422 | 0.3117 | 9 |
| Trichostatin A | GO:0006325 | chromatin organization | 23 | **\*0.0169** | 3 | 24 | **0.0173** | 1 |
| Tunicamycin | GO:0070085 | glycosylation | 1 | **\*0.0005** | 1 | 1 | **0.0005** | 1 |
| Tyrocidine B | GO:0071554 | cell wall organization or biogenesis | 5 | **\*0.0040** | 1 | 2 | **0.0019** | 1 |
|  |  | Num with significant rank |  | 22 |  |  | 21 |  |
|  |  | Num with significant rank and FDR < 25% |  | 16 |  |  | 0 |  |

Given that the main performance advantage of CG-TARGET occurred in the context of controlling the false discovery rate, we conclude that the issues with simple enrichment-based approaches primarily emerge not when predicting the most likely perturbed bioprocess for any single compound but when comparing the strength and significance of bioprocess predictions across compounds to prioritize compounds from a large-scale chemical-genetic interaction screen. The aforementioned rank-based analysis of 35 gold-standard compound-bioprocess annotations supports this assertion, as none of the 21 significantly-ranked annotations predicted by top-20 enrichment passed the high-confidence threshold (FDR ≤ 25%), while 16 of the 22 significantly-ranked annotations predicted by CG-TARGET did so (Table 2). This difference between CG-TARGET and enrichment-based methods likely emerges from the ability of weak chemical-genetic interaction profiles to generate strong, statistically significant predictions in the absence of methods (such as CG-TARGET) that account for general signals that arise upon treatment with bioactive compound – especially if these signals are amplified through their similarity to a large cluster of profiles in the genetic interaction network. Thus, the substantially superior ability of CG-TARGET to control the false discovery rate relative to the enrichment-based approach is a critical quality in the context of large-scale, systematic compound screens.

### Characterizing performance with respect to individual bioprocess terms

In addition to benchmarking CG-TARGET’s ability to prioritize gold-standard annotated bioprocesses for specific compounds, we also benchmarked its ability to prioritize compounds that perturb specific bioprocesses. Specifically, each GO term was evaluated based on the ranks of the predictions for the simulated chemical-genetic interaction profiles derived from genes annotated to that GO term. The 100 best-performing terms represented a diversity of bioprocesses related to the proteasome, glycolipid metabolism, DNA replication and repair, replication and division checkpoints, RNA splicing, microtubules, Golgi and vesicle transport, and chromatin state (Fig S2). In contrast, the 100 worst-performing terms were bioprocesses primarily related to carbohydrate, nucleotide, and coenzyme/cofactor metabolism, as well as the mitochondria, transmembrane transport, and protein synthesis and localization (Fig S3). The best-performing terms were also significantly smaller than the worst-performing ones (8 and 35 genes on average, respectively; rank-sum p-value < 2.2 × 10^−16^), which, given the fact that we would expect the power to increase with gene set size assuming the corresponding set was still functionally coherent, suggests that our method identifies functionally specific signal. Interestingly, the relatively poor performance of many metabolism-related bioprocess terms may result from the fact that the chemical-genetic and genetic interaction screens were both performed in relatively rich medium, precluding analysis of condition-specific phenotypes for genes only required for growth in minimal medium. While the set of best-performing terms did include a diverse range of bioprocesses, the possibility of “blind spots” should always be considered when interpreting the predictions made by CG-TARGET, as they may lead to false negative results that either exclude interesting compounds (e.g. those whose primary modes of action affect carbohydrate metabolism) or mask potential side effects of compounds whose primary modes of action are more easily observed by this method.

### Application of CG-TARGET to protein complexes refines functional specificity of mode-of-action predictions

The prediction of perturbed protein complexes offers the opportunity to enhance the specificity of GO biological process predictions (especially for overly-general bioprocess terms) and investigate functional space not accessible by bioprocess annotations. As such, we investigated the potential to expand the use of CG-TARGET to the prediction of perturbed protein complexes. When CG-TARGET was applied to predict protein complex targets for the RIKEN screen data, 714 compounds were identified with at least one high-confidence (FDR ≤ 25%) complex prediction, 604 of which also occurred in our original set of RIKEN compounds with high-confidence bioprocess predictions. Similar, but not completely overlapping, sets of genes (Jaccard index > 0.2) contributed to the top 5 of both bioprocess and protein complex predictions for more than one third of these compounds (219; 36%); this suggested that the two standards possessed both shared and complementary functional information that could be used to improve predictions.

We observed that protein complex predictions narrowed down less-specific bioprocess terms and enabled predictions in places where bioprocess annotations were sparser. To assess the ability to refine bioprocess prediction specificity, we mapped each protein complex to the childless bioprocess terms that completely encompassed them and looked for substantial improvements in prediction strength from the bioprocess to its protein complex “child.” We observed several instances in which bioprocess predictions with FDR > 25% (not high confidence) could be converted to high-confidence predictions by refining the bioprocess term to a constituent protein complex. For example, we saw substantial gains for the following bioprocess-to-complex combinations (sizes in parentheses): “mRNA polyadenylation” (bioprocess, not high confidence; size 8) to “mRNA cleavage factor matrix” (complex, high confidence; size 4); “cytoplasmic translation” (51) to “cytoplasmic ribosomal large subunit” (24); “vacuolar acidification” (14) to “H^+^-transporting ATPase, Golgi/vacuolar” (5); and “regulation of fungal-type cell wall organization” (8) to PKC pathway” (4) (Table S1). Importantly, 27 of the 110 compounds with high-confidence protein complex but not bioprocess predictions achieved their high-confidence status purely based on protein complex predictions that enhanced the specificity of a non-high-confidence, overlapping bioprocess prediction. Additionally, a separate set of 22 out of 110 compounds achieved high-confidence status based solely on predictions to protein complexes that did not strongly overlap with any bioprocesses (Jaccard < 0.2), demonstrating that the current set of protein complex annotations enabled predictions in functional space that was not well captured by a GO biological process term.

### Assessing the compatibility of chemical-genetic and genetic interaction profiles

Our evaluations of CG-TARGET support the premise of the method that genetic interaction profiles can be used as a tool to interpret chemical-genetic interaction profiles. However, we sought to better understand the extent to which these two types of profiles actually agree with one another, and if their systematic differences could shed light on the limits of the core assumption behind our method (i.e. that chemicals mimic the interaction profiles of their genetic targets). To investigate the compatibility of chemical-genetic and genetic interaction profiles, we quantified the contribution of individual gene mutants in the chemical-genetic interaction profiles to the prediction of individual bioprocesses. For a single compound and predicted bioprocess, these “importance scores” were obtained by 1) computing a mean genetic interaction profile across all *L*_2_-normalized query genetic interaction profiles that possessed an inner product of 2 or higher with the chemical-genetic interaction profile and mapped to the predicted bioprocess, and 2) computing the Hadamard product (elementwise multiplication) between this mean genetic interaction profile and the compound’s chemical-genetic interaction profile. Each score could have been positive, indicating agreement in the sign of chemical-genetic and genetic interactions for a gene mutant, or negative, indicating that the interactions did not agree for that gene mutant. As such, the importance scores summarized the concordance between chemical-genetic and genetic interaction profiles, conditioned on an individual compound and a perturbed bioprocess of interest.

We use the prediction of NPD4142, a compound from the RIKEN Natural Product Depository, to the “mRNA transport” bioprocess to illustrate how the overlap between chemical-genetic and genetic interactions led to bioprocess predictions (Fig 4A). A qualitative examination revealed that, indeed, NPD4142 possessed a pattern of chemical-genetic interactions similar to the genetic interactions for the query genes annotated to mRNA transport. More quantitatively and as expected, we observed that the contribution of each gene mutant to a bioprocess prediction depended on the strength of its chemical-genetic interaction with NPD4142 and the number and intensity of its genetic interactions with the mRNA transport query genes. Chemical-genetic interactions with mutants of *POM152, NUP133*, and *NUP188*, which encode components of the nuclear pore that facilitate import and export of molecules such as mRNA, were the most important, followed by interactions with mutants in the Lsm1-7-Pat1 complex, which is involved in the degradation of cytoplasmic mRNA.

**Figure 4.**
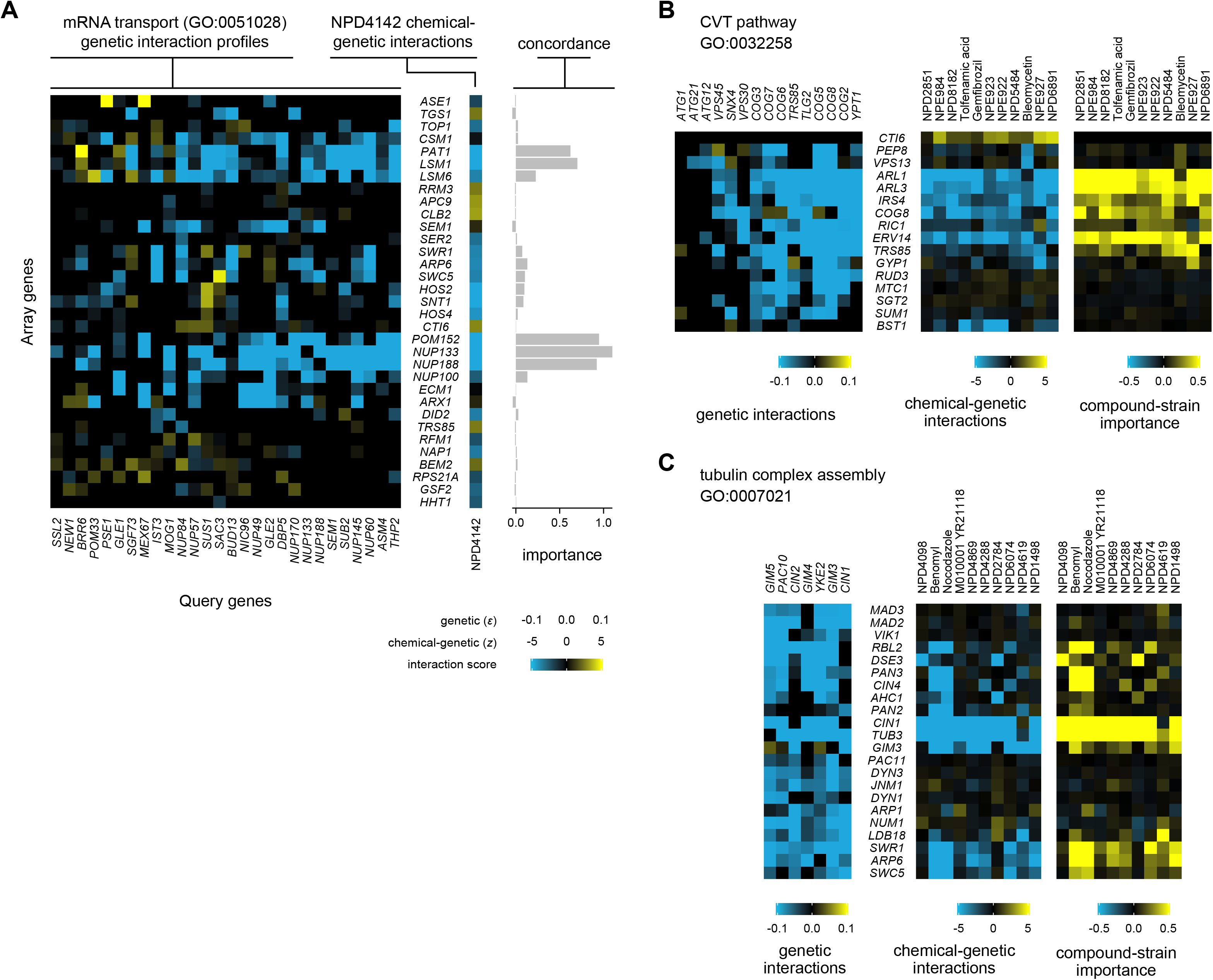
Detailed analysis of the contribution of individual gene mutants to biological process predictions. Each panel shows, for a bioprocess and either a compound (A) or a set of compounds (B-C) predicted to perturb that bioprocess, the subset of the respective chemical-genetic and *L*_2_-normalized genetic interaction profiles with signal. The importance profiles are the row-wise mean of the Hadamard product (elementwise multiplication) of each chemical-genetic interaction profile and the genetic interaction profiles for query genes with which it possessed an inner product of 2 or higher that are annotated to the GO term; they reflect the strength of each strain’s contribution to the bioprocess prediction. For all panels, a query gene from the genetic interaction network was selected if it contributed to the importance score calculation for any selected compound; query genes were ordered from left to right in ascending order of their inner products (or their average, for B-C) with the selected chemical-genetic interaction profile(s). Each strain (row) was included if it passed at least one of three criteria: 1) the magnitude of its mean genetic interaction score across the selected query genes exceeded 0.04; 2) the magnitude of its chemical-genetic interaction score (for B-C, the mean of such scores) exceeded 2.5; or 3) its importance score exceeded 0.1 (for B-C, the mean of such scores). (A) Schematic showing the prediction of the “mRNA transport” bioprocess (G0:0051028) for chemical compound NPD4142. (B) Schematic showing the prediction of “CVT pathway” (FDR < 1%) for compounds whose top prediction was to that term. (C) Schematic showing the prediction of “tubulin complex assembly” (FDR <1%).

Using this approach to assess the importance of individual mutants in the chemical-genetic profile, we globally analyzed the contribution of chemical-genetic interactions to each compound’s top bioprocess prediction (Fig 5). We performed this analysis twice: first, on all HCS compounds, and second, on a diverse subset of 130 compounds to correct for potential functional biases in the full set [11]. We present here the results from the 130-compound subset, although the results for the full set were qualitatively similar. For each compound, an average of 42% of its chemical-genetic interactions contributed to its top bioprocess prediction (chemical-genetic interaction cutoff ± 2.5, importance score cutoff +0.1) – a fraction that increased substantially (to 78%) when limiting the analysis to each compound’s strong interactions that contributed strongly (chemical-genetic interaction cutoff ± 5, importance score cutoff +0.5).

**Figure 5.**
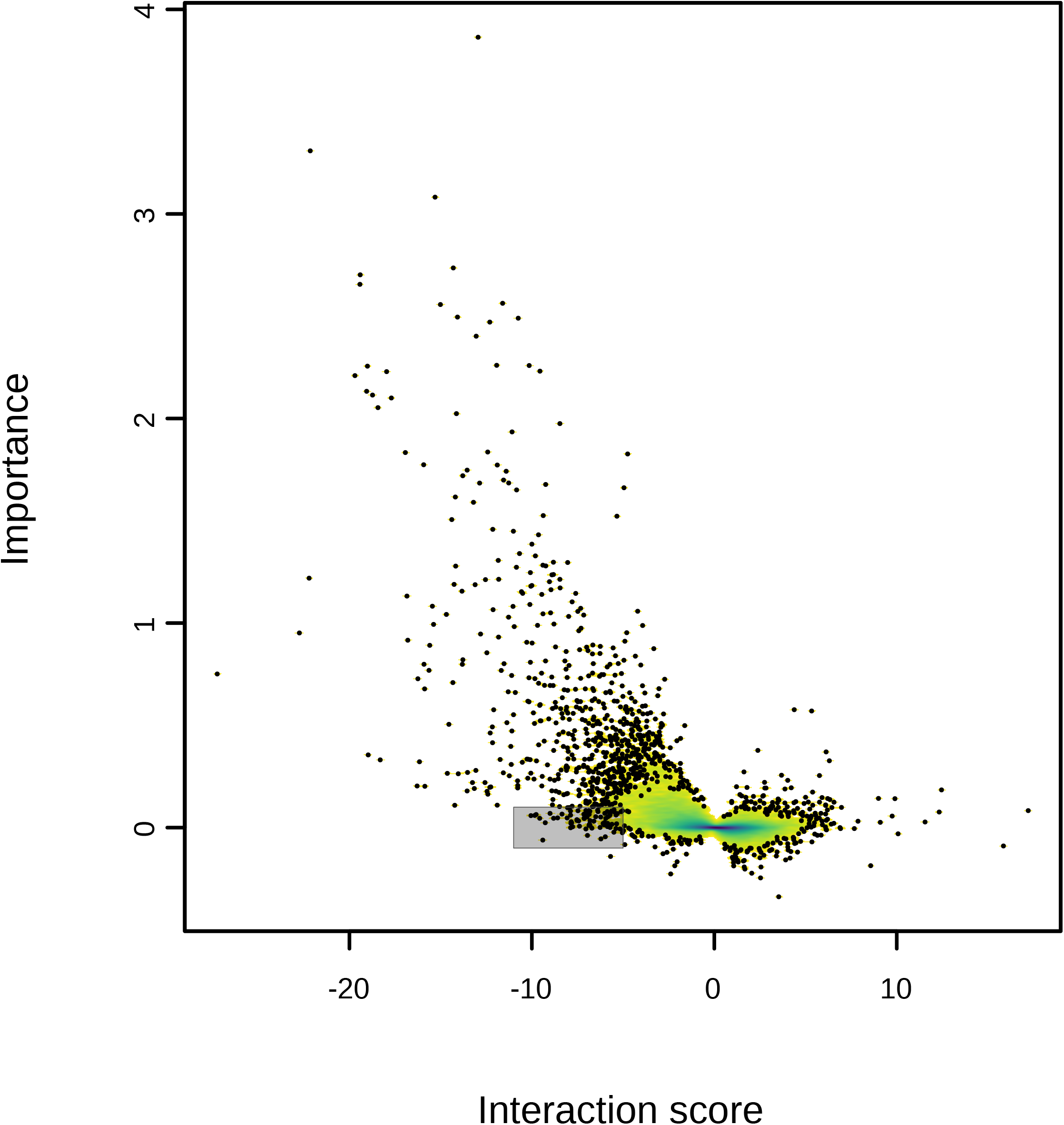
Global visualization of the contribution of chemical-genetic interactions to CG-TARGET bioprocess predictions. Chemical-genetic interaction profiles and their corresponding importance score profiles (see Fig 4 legend) were gathered for each of 130 diverse compounds from the high confidence set (FDR ≤ 25%) and their associated top bioprocess predictions. Importance is plotted as a function of chemical-genetic interaction score. One thousand points from the regions of lowest density (white) are plotted, with only density plotted in the remaining higher-density regions. Density increases in order of white, yellow, green, and violet. The shaded region highlights strains with strong negative (≤ −5) chemical-genetic interactions and no contribution (± 0.1) to a compound’s top bioprocess prediction.

Overall, we observed that more than one-third of chemical-genetic interactions (1112 / 3129) contributed to a top bioprocess prediction (chemical-genetic interaction cutoff ±2.5; importance score cutoff +0.1). Strikingly, negative chemical-genetic interactions much more frequently contributed to a bioprocess prediction: approximately one-half (1071 / 2112) of negative chemical-genetic interactions contributed as compared to only ~4% (41 / 1017) of positive chemical-genetic interactions at the same cutoff. Furthermore, we observed differences in how the signs within chemical-genetic and mean genetic interaction profiles could disagree with each other despite the global profile similarity that led to bioprocess prediction, with positive chemical-genetic interactions contributing negatively to bioprocess predictions (importance score cutoff < −0.1) over 10 times more frequently than negative interactions (1.9% vs. 0.14%). This trend of negative chemical-genetic interactions supporting strong bioprocess predictions was even more pronounced when restricting this analysis to strong interactions (chemical-genetic interaction cutoff ±5; importance score cutoff +0.5), where negative interactions comprised essentially the entire set of contributing chemical-genetic interactions (219 / 220, 99.5%). These observations were also supported by analyses in which we predicted perturbed bioprocesses using only negative or positive chemical-genetic interactions, finding that negative chemical-genetic interactions were the primary drivers of bioprocess predictions and overwhelmingly responsible for their accuracy [11]. We conclude that negative interactions in chemical-genetic interaction profiles contain the large majority of the functional information necessary to predict modes of action.

Negative chemical-genetic interactions also contained information specific to chemical perturbations. Specifically, we identified nine mutant strains that exhibited strong negative chemical genetic interactions (z-score < −5) yet were enriched for a lack of contribution (importance score < 0.1) to bioprocess predictions (hypergeometric test, Benjamini-Hochberg FDR ≤ 0.05; shaded region of Fig 5). Manual inspection of these mutants revealed connections to the high osmolarity glycol (HOG) pathway, cell polarity (cytoskeletal actin polarization, kinetochore and chromosome segregation), and other stress response mechanisms (Table S2). As the HOG pathway is important for the cellular response to high osmolarity and other stresses [18–20], and repolarization of the cytoskeleton is required for cells to adapt and continue dividing after stress [21,22], we hypothesize that many of these overrepresented mutants interact negatively with compounds due to an impaired ability to respond to external stress. This chemical perturbation-specific information may complement or even completely obscure the chemical-genetic signature of a compound’s primary mode of action, potentially complicating the interpretation of chemical-genetic interaction profiles using a genetic interaction network.

We compared the concordance of chemical-genetic and genetic interaction profiles across multiple compounds predicted to the same bioprocess, revealing that some bioprocesses were predicted by homogenous sets of chemical-genetic interaction profiles while others were much more heterogeneous despite their predicted targeting of the same bioprocess. For example, predictions made to the “CVT pathway” (FDR < 1%) depended almost entirely on a suite of strong negative chemical-genetic interactions with *ARL1, ARL3*, and *ERV13*, with contributions from *IRS4* and *COG8* (Fig 4B). This uniformity in the prediction of a bioprocess is contrasted by the diversity of profiles captured within “tubulin complex assembly” predictions (Fig 4C). Compounds with top predictions to this term could potentially be partitioned into three classes, divided according to strong contributions from: 1) *CIN1/TUB3, PAN3/CIN4*, and the SWR1 complex (known tubulin polymerization inhibitors Benomyl and Nocodazole); 2) *CIN1/TUB3* and *DSE2* (NPD4098 and NPD2784); or 3) only *CIN1/TUB3* (all remaining compounds except NPD4619). Interestingly, the structures of the compounds in each of the former two groups are distinct from those in the other groups, suggesting that the observed diversity in these compounds’ functional profiles is mechanistically derived from their structures.

### Experimental validation of compound-bioprocess predictions

**Phenotypic analysis of cell cycle progression**. The genes and pathways that govern the cell cycle are highly conserved throughout eukaryotes, enabling researchers to infer from yeast how cells in higher organisms integrate internal and external signals to decide when to divide [23]. As such, compounds that inhibit the progression of the cell cycle in yeast may enable a better understanding of the eukaryotic cell cycle or even form the basis for new therapeutic approaches for cancer, in which the cell division cycle is dysregulated [24,25]. We observed that compounds from the RIKEN Natural Product Depository were enriched for predictions to cell cycle-related bioprocesses [11], especially to the “mitotic spindle assembly checkpoint” that occurs at the beginning of M phase. After manual inspection of these compounds’ chemical-genetic interaction profiles, we selected 17 to test if our predictions validated experimentally. Specifically, we looked for increases in the percentage of cells in the G2 phase of the cell cycle (via fluorescence-activated cell sorting) and two budding phenotypes (bud size and % cells with large buds) for yeast treated with compound, together indicative of arrest at the G2/M checkpoint of the cell cycle (Fig 6A-C). Indeed, 6 of the 17 selected compounds induced increases in all phenotypes, while zero out of 10 bioactive control compounds (with high-confidence predictions to bioprocesses not related to cell cycle signaling and progression) induced increases in any of these phenotypes (p < 0.05, one-sided Fisher exact test). As compounds can activate the G2/M checkpoint in multiple ways (e.g. induction of DNA damage, inhibition of chromosome segregation), the set of compounds with spindle assembly checkpoint predictions can serve as a resource for studying the diversity of mechanisms by which cell cycle progression is arrested at this checkpoint and which of these may have therapeutic potential. In addition to our study of G2/M checkpoint-activating compounds, we also selected two compounds with high-confidence predictions to the term “cell-cycle phase” (mutually exclusive with mitotic spindle assembly checkpoint), one of which (NPD7834) was observed to arrest cells in G1 phase (Fig 6A-C).

**Figure 6.**
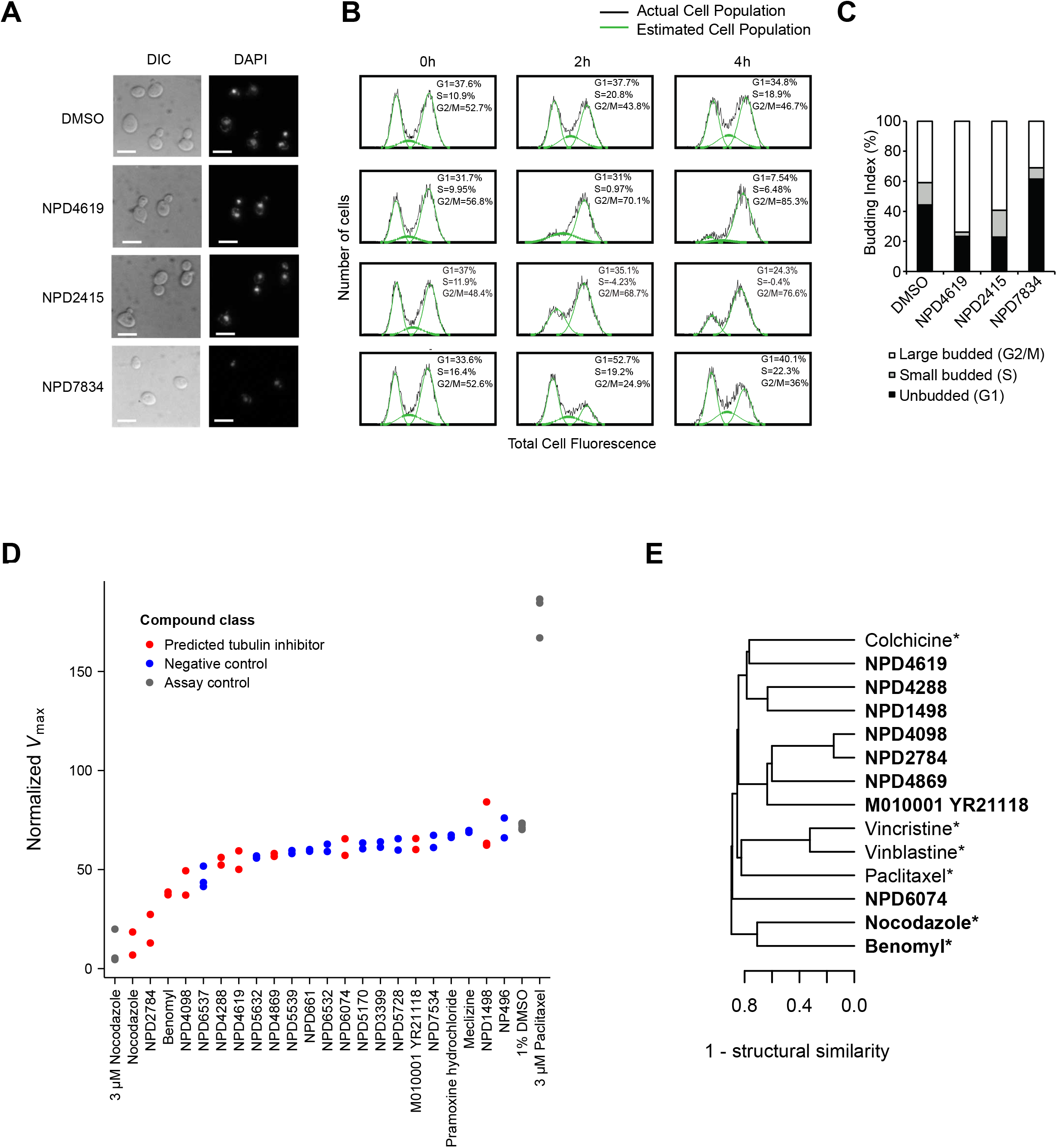
*In vivo* and *in vitro* experimental validations of biological process predictions. (A,B,C) Phenotypic validation of cell cycle-related predictions, performed on drug-hypersensitive yeast treated with solvent control (DMSO) or compounds predicted to perturb the cell cycle. (A) Differential interference contrast microscopy (DIC) and fluorescence upon DAPI staining showing bud size and DNA localization, respectively, after compound treatment. The scale bar represents a distance of 5 μm. (B) FACS analysis of cell populations in different cell cycle phases at 0, 2, and 4 hours after compound treatment. The green curve overlay represents the estimated cell population in G1, S and G2/M phases. (C) Budding index percentages induced by treatment with compound or solvent control. (D) *In vitro* inhibition of tubulin polymerization by compounds predicted to perturb “tubulin complex assembly” (FDR < 1%; red) compared to randomly-selected negative control compounds with high-confidence predictions to bioprocesses not related to chromosome segregation, kinetochore, spindle assembly, and microtubules (blue). *V*_max_ values reflecting the maximum rate of tubulin polymerization for each compound from independent replicate experiments are plotted. Assay positive and negative control compounds are colored grey. (E) Structural similarity-based hierarchical clustering of compounds tested in (D). Single linkage was used in combination with (1 – structural similarity) as the distance metric; as such, the structural similarity of the two most similar compounds at each junction can be inferred directly from the dendrogram. Compounds predicted to perturb “tubulin complex assembly” (FDR < 1%) are in bold, and known microtubule-perturbing agents are marked with an asterisk. Structural similarity was calculated as the Braun-Blanquet similarity coefficient on all-shortest-path chemical fingerprints of length 8 (see Materials and Methods).

**Inhibition of tubulin polymerization**. Compounds that disrupt microtubules are useful for studying cell organization and division and remain promising candidates as antitumor agents [26–28]. As such, we focused on all compounds with the strongest predictions to “tubulin complex assembly” (FDR < 1%) and tested them for activity in an *in vitro*, mammalian (porcine) tubulin polymerization assay (Fig 6D). Like the previous validation experiment, a negative control set of compounds was selected at random to contain high-confidence compounds (bioprocess predictions with FDR ≤ 25%) whose predictions were not related to microtubule assembly or related bioprocesses. We observed that the novel compound NPD2784 strongly inhibited tubulin polymerization, nearly as well as the drug nocodazole and more strongly than the microtubule probe benomyl. In addition, the entire set of compounds predicted to perturb tubulin complex assembly showed significantly increased inhibition of tubulin polymerization when compared to the negative control compounds (p < 0.006, Wilcoxon rank-sum test). Strikingly, all previously-uncharacterized members of this set would not have been discovered using a structure similarity-based approach, as the highest structural similarity between any NPD compound and six compounds representative of major classes of microtubule-perturbing agents did not exceed 0.25 (Fig 6E) [29]. However, we did observe that structural similarity was predictive of the top 20% of chemical-genetic profile similarities among the compounds selected for validation (AUPR = 0.43 vs. 0.2 for a random classifier), suggesting that their slight differences in function inside the cell are influenced by their structures and that further exploration of compounds with similar structures may yield even more tubulin polymerization inhibitors. With this experimental validation, we have demonstrated the ability of CG-TARGET, and a genetic interaction network in general, to capture a shared mode of action across diverse compounds that can be biochemically-validated. Furthermore, we note that this validation was achieved with a mammalian tubulin assay, demonstrating the power of yeast chemical genomics coupled with CG-TARGET to predict modes of action that translate broadly to other species, including mammalian systems.

## Discussion

The scaling of chemical-genetic interaction screens from tens or hundreds of compounds to tens of thousands of compounds has provided the opportunity, and the necessity, to develop better methods for interpreting the interaction profiles and prioritizing high-confidence compounds. We developed a method, CG-TARGET, to address this need and used it to predict perturbed biological processes for the nearly 14,000 compounds interrogated in our recent high-throughput chemical-genetic interaction screen [11]. CG-TARGET demonstrated the ability to recapitulate known compound function while controlling the false discovery rate, enabling high-confidence mode-of-action prediction for 1522 largely uncharacterized compounds [11], which we prioritized for further study. Further investigation of the profiles from these high-confidence compounds revealed broad compatibility between chemical-genetic and genetic interaction profiles, the overwhelming basis of which was contributed by negative chemical-genetic interactions. Some interesting exceptions to this compatibility were observed for genes that may reduce the ability of compounds to deal with external stress. We experimentally confirmed the accuracy of our predictions for two different classes of previously uncharacterized compounds – tubulin polymerization inhibitors and mitotic checkpoint inhibitors – and demonstrated the ability of CG-TARGET to predict activity against a conserved mammalian target. In addition to these findings, the predictions made using CG-TARGET were experimentally validated on a large scale for 67 compounds in an orthogonal cell cycle assay and revealed insights into the distribution of functions perturbed by compounds in large compound libraries, which is described in the companion paper [11].

In high-throughput chemical screens, it is important to prioritize the compounds most likely to demonstrate desired biological activity in further studies. While CG-TARGET and a baseline, enrichment-based approach achieved similar performance in ranking gold-standard bioprocess annotations for simulated chemical-genetic interaction profiles and compounds with known modes of action, CG-TARGET outperformed the baseline approach with regard to controlling the false discovery rate, discovering two orders of magnitude more compounds at a false discovery rate of 25%. As a result, CG-TARGET was substantially better than the baseline approach at accurately annotating, with high confidence, compounds with known modes of action. The fact that our genetic interaction-based predictions were both accurate and achieved appropriate control of the false discovery rate is important, as the global genetic interaction network provides a much more comprehensive and unbiased resource than the limited set of gold standard compounds for predicting bioprocesses perturbed by compounds. In addition, predicting compound function at the bioprocess level allowed functional characterization of compounds whose effects in cells did not occur via direct action on protein targets (e.g. damaging DNA or disrupting cell membranes,), which would have been impossible with a method based purely on comparing chemical-genetic and genetic interaction profiles.

While we demonstrated the ability to predict perturbed bioprocesses for compounds and prioritize the highest-confidence predictions, many further steps are required to identify lead compounds and ultimately develop molecular probes or pharmaceutical agents. Perturbing a biological process does not necessarily require perturbing a specific protein target, and as such, further refinements to our methods are needed to identify specific molecular targets (i.e. proteins) and prioritize the compounds most likely to perturb a small number of defined targets in the cell. We envision the use of multiple functional standards with CG-TARGET, such as biological processes and protein complexes as demonstrated here, to improve our ability to predict compound mode of action at different levels of resolution and predict the compounds that exert specific versus general effects in the cell. Different modes of chemical-genetic interaction screening can provide support in this endeavor, as heterozygous diploid mutant strains, gene overexpression strains, and/or spontaneous compound-resistant mutants can provide evidence for the direct, essential cellular target(s) of a compound [1,7]. Regardless of the limitations in predicting precise molecular targets, information about the bioprocesses perturbed by an entire library would be useful in selecting the compounds most amenable to activity optimization and off-target effect minimization in the development of a pharmaceutical agent or molecular probe.

The approach described here can be translated to work in other species for which obtaining functional information on compounds would be useful. For example, genome-wide deletion collections have been developed for *Escherichia coli* [30] and *Schizosaccharomyces pombe* [31] and used to perform chemical-genetic interaction screens [32,33] as well as genetic interaction mapping [34–37]. Such efforts are even underway in human cell lines, enabled by genome-wide CRISPR screens [38–41]. Furthermore, future efforts to interpret chemical-genetic interaction profiles in a new species need not wait for the completion of a comprehensive, all-by-all genetic interaction network as exists in *S. cerevisiae*, as our work highlights the ability of a diagnostic set of gene mutants to capture functional information and predict perturbed biological processes. From the discovery of urgently-needed antibacterial or antifungal agents, to the treatment of orphan diseases or a better understanding of drug and chemical toxicity, the combination of chemical-genetic and genetic interactions in a high-throughput format, with appropriate analysis tools, offers a means to achieve these goals via the discovery of new compounds with previously uncharacterized modes of action.

## Materials and Methods

### Datasets

**Chemical-genetic interaction data**. Chemical-genetic interaction profiles were obtained from a recent study [11], in which nearly 14,000 compounds were screened for chemical-genetic interactions across ~300 haploid yeast gene deletion strains. The chemical-genetic interaction profiles consisted of two sub-datasets: 1) the “RIKEN” dataset, containing chemical-genetic interaction profiles spanning 289 deletion strains for 8418 compounds from the RIKEN Natural Product Depository [12] and 5724 negative experimental controls (solvent control, DMSO); and 2) the “NCI/NIH/GSK” dataset, containing chemical-genetic interactions spanning 282 deletion strains for 3565 compounds from the NCI Open Chemical Repository, the NIH Clinical Collection, and the GSK kinase inhibitor collection [13], as well as 2128 negative experimental control profiles. The solvent control profiles consisted of biological and technical replicate profiles.

**Genetic interaction data**. The genetic interaction dataset was obtained from a recently assembled *S. cerevisiae* genetic interaction map [5,9]; it was filtered to contain quantitative fitness observations for double mutants obtained upon crossing 1505 high-signal query gene mutants into an array of 3827 array gene mutants. The procedure for selecting the 1505 high-signal query genes out of the larger pool of 4382 is described in [11]. Briefly, each query profile was required to possess at least 40 significant genetic interactions, a sum of cosine similarity scores with all other query profiles greater than 2, and a sum of inner products with all other query profiles greater than 2. The final genetic interaction dataset used in this study was filtered to contain only array strains present in the chemical-genetic interaction datasets.

**GO Biological Processes and protein complexes**. A subset of terms from the “biological process” ontology within the Gene Ontology annotations [17] were used as the bioprocesses. Query genes from the *S. cerevisiae* genetic interaction dataset were mapped to biological process terms using annotations from the *Saccharomyces cerevisiae* Genome Database [16]. Both gene ontology and *S. cerevisiae* annotations were downloaded on September 12, 2013 from their respective databases via Bioconductor in R [42]. Terms were propagated using “is_a” relationships, such that each gene was also annotated to all parents of its direct biological process annotations. The final set of bioprocesses consisted of the terms with 4 – 200 gene annotations from the set of 1505 high-signal query genes in the genetic interaction dataset.

Protein complex annotations were obtained from [9]. Complexes with 3 or more genes annotated to them were used as the input biological processes for CG-TARGET-based protein complex predictions.

**Gold-standard compound-process annotations**. Biological processes were assigned to 35 primarily antifungal compounds with chemical-genetic interaction profiles in the RIKEN dataset, based on known information about their modes of action. Bioprocess terms were selected to be specific to the compounds’ modes of action where applicable.

### Predicting perturbed bioprocesses from chemical-genetic interaction profiles

Our method to predict biological processes perturbed by compounds is briefly summarized in the recent study from which the chemical-genetic interaction profiles were obtained [11], and is more formally described here. Fig S4 provides a schematic representation of the method.

**Notation**. We first clarify here a few uses of mathematical notation that simplify the explanation of the methods. First, the i^th^ row and column vectors of a matrix *A* are denoted as *A_i,*_* and *A_*,i_*, respectively. Second, the Iverson bracket is used to convert logical propositions into values of 1 or 0, depending on if the logical proposition is true or false, respectively. This is used to simplify expressions for counting the number of elements in a vector that meet given criteria. Specifically, for a logical proposition *L*, the definition of the Iverson bracket is:

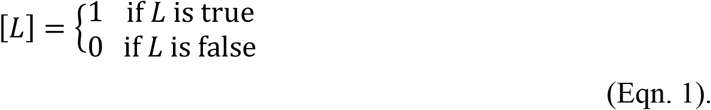

**Data representation and overview of procedure**. CG-TARGET requires chemical-genetic interaction profiles, genetic interaction profiles, and a mapping from genes to biological processes, all of which will be represented as matrices here (illustrated in Fig S4, along with example matrix dimensions and a graphical description of the bioprocess prediction procedure). For chemical-genetic interaction matrices, let us consider an *n_m_* × *n_α_* matrix of compound treatment profiles *C_α_*, an *n_m_* × *n_β_* matrix of negative experimental control profiles *C_β_*, and an *n_m_* × *n_γ_* matrix of resampled profiles *C_γ_*, where *n_m_* is the number of mutant strains in each chemical-genetic interaction profile, *n_α_* is the number of profiles derived from treatment with compound, *n_α_* is the number of profiles derived from negative experimental controls, and *n_γ_* is the number of chemical-genetic interaction profiles resampled from *C_α_*. The matrix *G* of genetic interaction profiles is *n_m_* × *n_q_* and the binary matrix *B* of gene to bioprocess mappings is *n_q_* × *n_p_*, where *n_m_* is the number of mutant strains in the chemical-genetic interaction and genetic interaction profiles, *n_q_* is the number of genetic interaction profiles, and *n_p_* is the number of bioprocesses in *B* annotated from the *n_q_* genetic interaction profiles in *G*.

To predict perturbed biological processes, chemical-genetic interaction matrices for each profile type *a* ∈ {*α*, *β, γ*} are first converted to matrices of compound-gene similarity scores and then to matrices containing the sums of these compound-gene similarity scores for each compound-process pair. Three different z-score/p-value matrix pairs are then computed for each profile type *a*, two of which are derived from the control chemical-genetic interaction profile types *b* ∈ {*β, γ*} (“control-derived” z-scores/p-values) and one of which is derived by randomizing the scores within each compound’s vector of compound-gene similarity scores (“within-compound” z-scores/p-values, denoted as *δ*). The z-score and p-value matrices across all scoring approaches *c* ∈ {*β, y, δ*} are then combined into a final z-score/p-value matrix pair for each profile type *a*. The false discovery rate is estimated by comparing the rate of prediction for the treatment profiles *α* against that of the control profiles *b* ∈ {*β, γ*} across a range of p-value thresholds. For the comparison of CG-TARGET to an enrichment-based approach, one enrichment factor/p-value matrix pair replaces the final z-score/p-value matrix pair for each profile type *a*, with the same false discovery rate calculations occurring afterward.

**Resampled chemical-genetic interaction profiles**. An *n_m_* × *n_γ_* matrix of resampled chemical-genetic interaction profiles *C_γ_* is constructed such that interaction scores for each gene are sampled randomly with replacement across the chemical-genetic interaction profiles. Assuming that rand(*x*) is a function to randomly sample one value from the set of integers *x* in a uniformly random fashion, and {1..*n_α_*} is the set of integers between and including 1 and *n_α_*, the interaction score for the *i*^th^ mutant in the *j*^th^ resampled profile is denoted by:

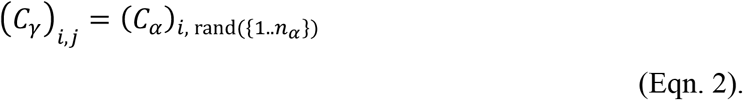

**Mapping the similarity between chemical-genetic and genetic interaction profiles onto biological processes**. Scores reflecting the concordance between chemical-genetic and genetic interaction profiles were derived by taking the inner product between each chemical-genetic interaction profile and each *L*_2_-normalized genetic interaction profile. As such, a column-normalized genetic interaction matrix *G'* is constructed from the genetic interaction matrix *G* by:

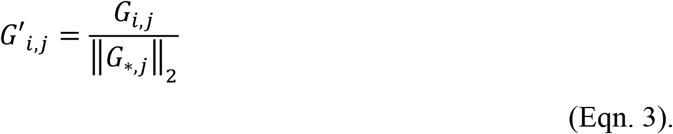

Matrices *S_α_* (*n_α_* × *n_q_*)*, S_β_* (*n_β_* × *n_q_*), and *S_R_* (*n* × *n_q_*), containing the similarity scores between the genetic interaction profiles and the profiles from each compound-treated, negative experimental control, and resampled condition, respectively, are then generated as denoted by (where the superscript *^T^* indicates the matrix transpose):

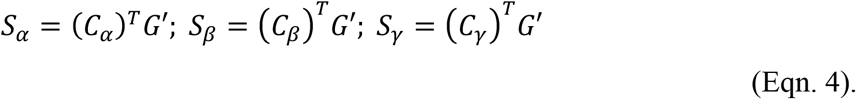

To map these similarity scores onto biological processes, the inner product is taken between each row vector of compound-gene similarity scores (from *S_α_, S_β_*, and *S_γ_*) and the column vector of binary gene annotations from each bioprocess in matrix B. This generates matrices *X_α_* (*n_α_* × *n_p_*)*, Χ_β_* (*η_β_* × *n_p_*), and *X_γ_* (*n_γ_* × *n_p_*) that contain the sum of gene similarity scores within each biological process for each compound treatment, negative experimental control, and resampled condition, respectively. These matrices are denoted by:

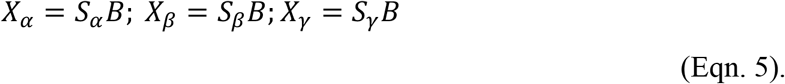

**Computing biological process predictions with CG-TARGET**. Once the compound-gene similarity scores are mapped onto biological processes and summed into compound-process scores, we compute z-score matrices 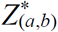 and empirical p-value matrices *P*_*z\**(*a, b*)_, where *a* denotes the type of profile we are predicting bioprocesses for and *b* denotes the type of control distribution used to compute the z-scores and p-values. For two of the values of *b* (*α* and *β*), these scores are “control-derived,” as we compare each compound-process score (*X_a_*)*_i,j_* to the distribution of control profile compound-process scores (*X_b_*)*_*,j_* within the respective *j*^th^ bioprocess. For the remaining value of *b* (*δ*), we refer to these scores as “within-compound,” as we compare the *i*^th^ compound’s average compound-gene similarity score within genes annotated to the *j*^th^ bioprocess (*X_a_*)*_ij/dj_* (where *d_j_* is the size of the *j*^th^ bioprocess) to the distribution of compound-gene similarity scores (*S_a_*)*_i,*_* for the *i*^th^ compound.

The computation of each control-derived z-score requires an estimate of the mean and standard deviation of the compound-process scores within each bioprocess for both the negative experimental control and resampled profiles. The length *n_p_* mean vector *u_b_* and standard deviation vector *v_b_* for each control profile type *b* ∈ {*β, γ*} are thus defined as:

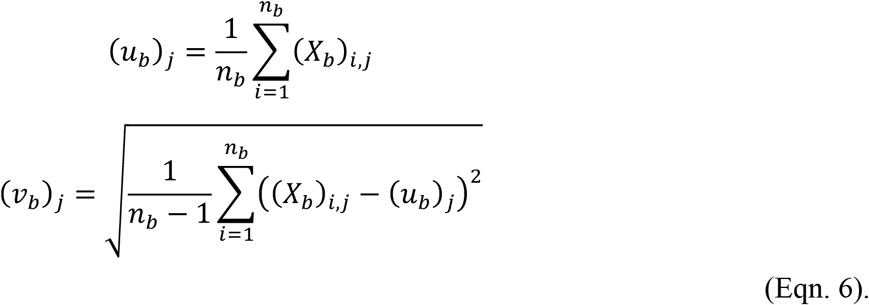

Z-score matrices derived using both types of control profile are computed for all compound treatment, negative experimental control, and resampled profile conditions, yielding six z-score matrices. These matrices, one for each combination of profile type *a* ∈ {*α*, *β, γ*} and control profile type *b* ∈ {*β, γ*}, are defined as:

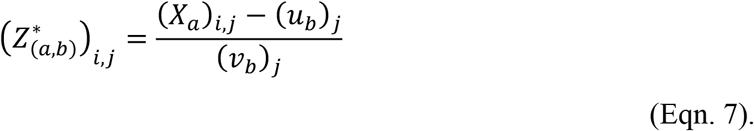

The control-derived p-values are computed by counting the number of times that a compound-process score (*X_a_*)*_i,j_* for the *i*^th^ compound and *j*^th^ bioprocess is less than the corresponding control-derived compound-process scores (*X_b_*)*_*,j_*. Again, this yields six p-value matrices, one for each combination of profile type a ∈ {α, β, γ} and control profile type b ∈ {β, γ}, which are given by:

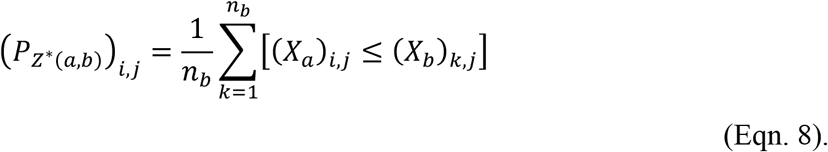

The within-compound z-score is computed for each pair of *i*^th^ compound and *j*^th^ bioprocess by comparing the mean of the *i*^th^ compound’s similarity scores with genes in the bioprocess to the mean and standard deviation of the *i*^th^ compound’s similarity scores across all genes. To perform this calculation, length *n_a_* mean and standard deviation vectors *w_a_* and *y_a_*, respectively, are generated, as well as a length *n_p_* vector *d* that contains the number of genes annotated to each bioprocess in *B*. 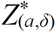 refers to the matrix of z-scores for each profile of type *a* ∈ {*α*, *β, γ*} computed using the within-compound z-score approach (represented by *δ*) and given by:

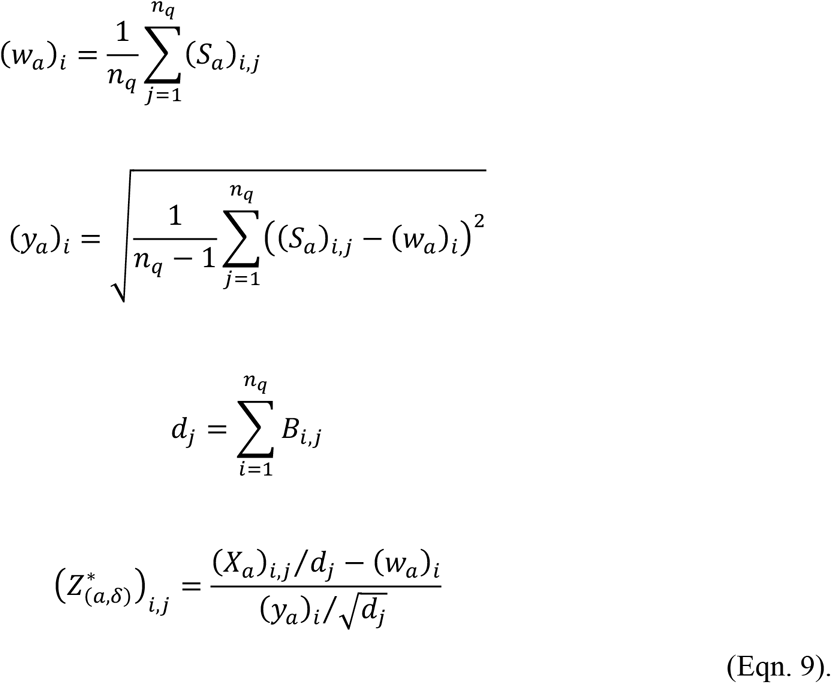

For each compound-process pair, the within-compound empirical p-value is computed for each profile type *a* ∈ {*α*, *β, γ*} by randomly permuting the compound’s compound-gene similarity scores, re-computing within-compound z-scores, and counting the number of times that the z-scores derived from randomly-permuted compound-gene similarity scores are greater than the observed compound-process z-score. This calculation conveniently reduces to a comparison of the sum of observed vs. permuted compound-gene similarity scores for genes in the respective bioprocess, as the number of genes that map to the bioprocess (*d_j_*) and the mean ((*w_a_*)*_i_*) and standard deviation ((*y_a_*)*_i_*) of compound-gene similarity scores do not change upon permutation of the compound-gene similarity scores. Permuted matrices of compound-gene similarity scores are denoted by *^k^S_a_*, which represents, for profile type *a*, the *k*^th^ row-wise permutation of the compound-gene similarity score matrix. Each resulting matrix that contains the sums of compound-gene similarity scores for all compound-process pairs with respect to random permutation *k* is denoted by *^k^X_a_*. Across *n_l_* permutations, the within-compound empirical p-value for each profile type *a* ∈ {*α*, *β, γ*} (within-compound p-value signified by subscript *δ*) is denoted by:

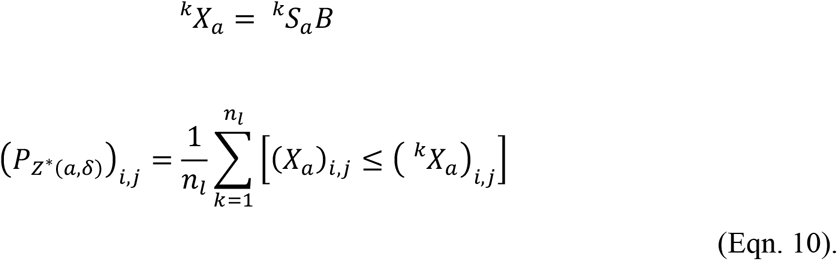

Ultimately, the different p-values and z-scores for each compound-process pair are combined into one p-value and z-score for that pair. These scores are combined such that the largest (least significant) p-value is chosen along with its associated z-score. If multiple p-values tie for the largest value, then the one with the smallest associated z-score is chosen. As such, the resulting combination of p-value and z-score represents the most conservative estimate of the strength and significance of the prediction from compound to perturbed biological process.

To combine the p-values and z-scores, a matrix *Psource_a_* for each profile type *a* ∈ {α, β, γ} is first created to determine, for each compound-process pair, which p-value and z-score matrices will contribute the final p-value and z-score. For each z-score/p-value scoring approach *c* ∈ { *β*, *γ, δ*}, each entry of this matrix is denoted by:

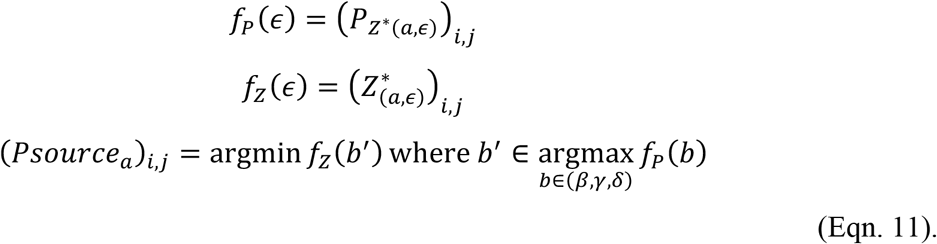

The resulting final p-value and z-score matrices for each profile type *a* ∈ (*α*, *β*, *γ*) are then:

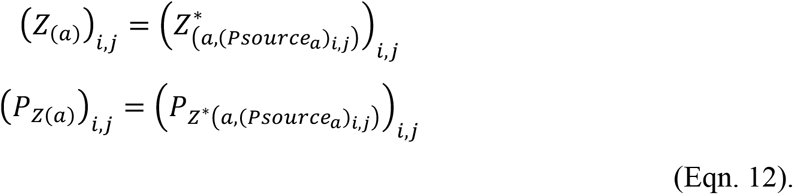

**Computing biological process enrichments**. An enrichment-based method for predicting biological processes perturbed by compounds was also implemented to provide an appropriate baseline for assessing the performance of CG-TARGET. This enrichment-based method computes biological process enrichment within the genes that contribute the top *n* out of *n_q_* compound-gene similarity scores for each compound (from each compound-gene similarity score matrix *X_a_* for profile types *a* ∈ {*α*, *β*, *γ*}. Ultimately, two sets of matrices are computed, *E_(a,n_*_)_ and *P_E(a,n_*_)_, which respectively contain the enrichment factor and hypergeometric p-value for each compound and biological process pair. Enrichments were computed for *n* ∈ {10, 20, 50, 100, 200, 300, 400, 600, 800}.

First, a binary matrix 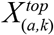 is derived from the matrix of compound-gene similarity scores *X_a_*, such that in each row, the positions corresponding to the top *n* scores are set to 1 and the remaining positions are set to 0. This is denoted as:

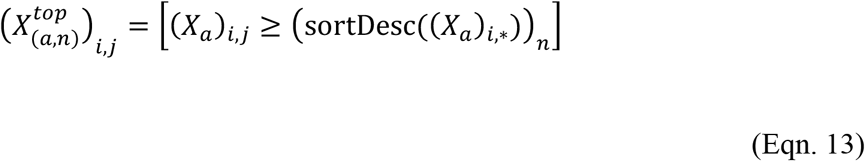
where (*X_a_*)*_i,*_* is the *i*^th^ row vector of matrix *X_a_* and sortDesc(*x*) is a function that returns the values in a vector x sorted in descending order. The final enrichment factor and p-value matrices are then computed as:

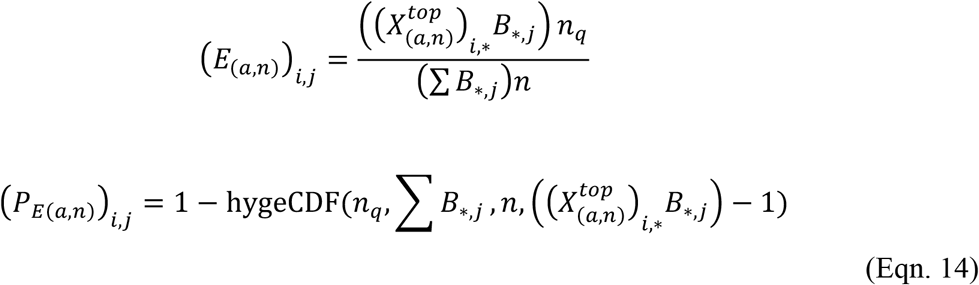
where *B_*,j_* is the column vector of the binary bioprocess matrix *B* containing gene annotations for the *j*^th^ bioprocess, 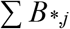 is the number of genes annotated to the *j*^th^ bioprocess, and hygeCDF(*N*, *K, n, k*) is the cumulative hypergeometric distribution given a population size of *N* with *K* success states and *n* draws with *k* observed successes.

**Estimating the false discovery rate**. The false discovery rates of the compound-process predictions are estimated by comparing, using the entire range of observed p-values as thresholds, the number of compounds with at least one bioprocess prediction against the number of experimental controls and resampled profiles with at least one bioprocess prediction. We compute a false discovery rate matrix *FDR_b_* for the treatment profiles *α* against each control profile type *b* ∈ {*β*, *γ*}. This *FDR_b_* matrix is individually computed for the CG-TARGET-based compound-process predictions as well as for each version of the enrichment-based compound-process predictions (using the p-value matrices *P_Z_*_(_*_a_*_)_ and *P_E_*_(_*_a,n_*_)_); for simplicity, we do not change the notation of *FDR_b_* to reflect if the false discovery rate values were computed on the output from CG-TARGET or our baseline enrichment-based approach.

The first step in computing the false discovery rate is obtaining length *n_α_* vectors *ptop_a_* that contain the smallest p-value within each profile’s bioprocess predictions, for each profile type *a* ∈ {*α*, *β, γ*}. Additionally, the union of all observed p-values *p_all_* defines the universe of p-values for which corresponding false discovery rates will be computed. Given p-value matrices *P_a_* (*P_Z_*_(_*_a_*_)_ or *P_E_*_(_*_a,n_*_)_ for one value of *n*) and a function sortAsc() that returns the input values sorted in ascending order, the vectors *ptop_a_* and *p_all_* are given by:

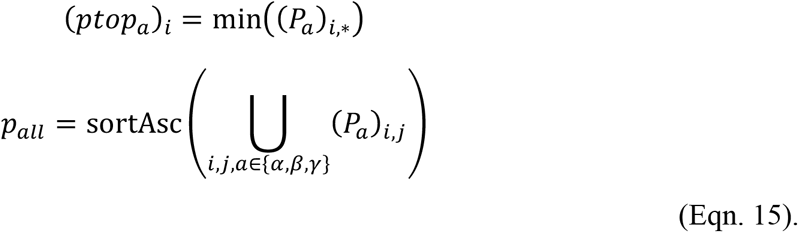

We then compute a mapping from each observed p-value to its corresponding false discovery rate, with mappings generated with respect to each control profile type *b* ∈ {*β, γ*}. First, a vector of false discovery rates 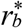 is computed, each value corresponding to a p-value threshold in *p_all_*, by dividing the fraction of treatment profiles with one or more bioprocess predictions that pass the threshold by the fraction of control profiles that also pass the threshold. As the p-values in the vector *p_all_* are monotonically increasing, it is desirable for the false discovery rate to increase monotonically with the p-value. However, it is possible for the false discovery rate to decrease as p-value increases (if the fraction of treatment profiles passing the threshold increases faster than the fraction of control profiles passing the threshold), and thus we adjust each false discovery rate value in the vector 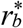 to be the minimum of its current value or any value at a larger index to generate a new vector *r_b_* (similar to the Benjamini-Hochberg procedure [43]). The final p-value to false discovery rate mappings can be written as a function of the p-value *p*, with the procedure to generate these mappings given by:

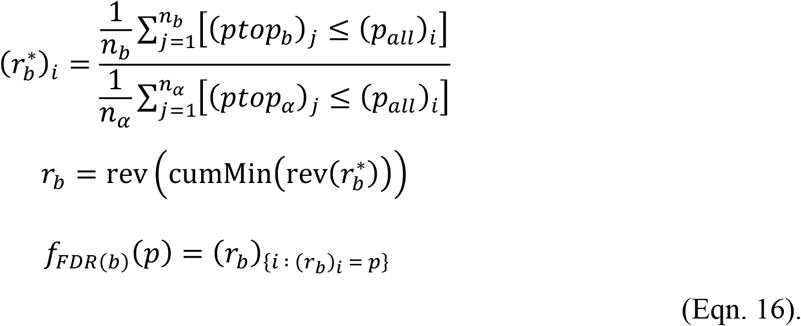

Given this mapping of p-value to false discovery rate, the resulting matrices of false discovery rates with respect to control profile types *b* ∈ {*β, γ*} are given by:

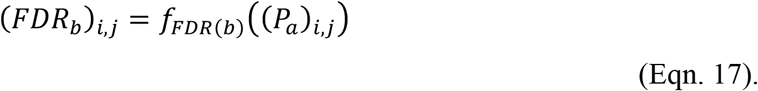

### Computational evaluation of bioprocess predictions

**Performance on simulated chemical-genetic interaction profiles**. We generated a set of simulated chemical-genetic interaction profiles derived from genetic interaction profiles [11]. Each simulated chemical-genetic interaction profile was a query genetic interaction profile augmented with noise sampled from a Gaussian distribution with a mean of 0 and a variance for each array gene twice that of the same array gene in the genetic interaction dataset. Three simulated profiles were generated based on each query gene, resulting in 4515 total profiles. Because each simulated chemical-genetic interaction profile was derived from a query genetic interaction profile, it inherited the gold-standard bioprocess annotations from its parent genetic interaction profile in subsequent benchmarking efforts.

We then used CG-TARGET and each top-*n* enrichment method to predict perturbed bioprocesses for this set of 4515 simulated chemicals x 289 deletion mutants. For each simulated chemical, its top bioprocess prediction was compared to the set of inherited gold-standard bioprocess annotations, counting as a true positive if the top prediction matched an existing annotation and a false positive if it did not. Precision-recall curves were then generated by sorting the list of each simulated chemical’s top bioprocess predictions (p-value ascending, z-score or enrichment factor descending) and computing the precision (true positives / (true positives + false positives)) and recall (true positives) at each point in this list.

**Performance on gold-standard compound-bioprocess annotations**. The predicted perturbed bioprocesses for each of the gold-standard compounds were sorted, first in ascending order by their p-value and then descending order by their z-score (for CG-TARGET) or enrichment factor (top-*n* enrichment), and the rank of each compound’s gold-standard bioprocess annotation was recorded. To assess the significance of each rank, each pair of p-value and z-score was randomly assigned to a new bioprocess (without replacement), the lists re-ordered, and the ranks of each compound’s target bioprocess re-computed. The empirical p-value for each gold-standard compound-process pair was computed as the number of times the rank from the shuffled bioprocesses achieved the same or better rank as the observed rank, divided by the number of randomizations. These randomizations were also used as a baseline against which to compare the number of compounds (out of 35) that achieved a given rank, as seen in Figs 3 and S1; the displayed ribbons were generated by calculating, for each rank, the relevant percentiles on the distribution of compounds with randomized predictions that achieved that rank. The “effective rank” of a compound’s gold-standard bioprocess annotation was determined as the minimum rank of any bioprocess term with which it possessed sufficient gene annotation similarity (overlap index ≥ 0.4, where the overlap index of two sets is defined as the size of the intersection divided by the size of the smaller set).

**Characterizing performance with respect to individual bioprocess terms**. For each propagated GO biological process term used for bioprocess prediction, we gathered all predictions to that term across the 4515 simulated chemical-genetic interaction profiles and sorted the predictions in ascending order by p-value and then in descending order by z-score. The area under the precision-recall curve (AUPR) was calculated across this sorted list of simulated compounds, with a true positive defined as the occurrence of a simulated compound that was annotated to the bioprocess (via the simulated compound’s parent gene). To obtain the final evaluation statistic for each GO term, this AUPR was divided by the AUPR of a random classifier, which is equal to the number of true positives divided by the total number of simulated compounds.

### Assessing the compatibility of chemical-genetic and genetic interaction profiles

**Analysis of bioprocess prediction drivers in chemical-genetic interaction data**. Given a compound and a predicted bioprocess, a profile of “importance scores” describes the contribution of each gene mutant to that compound’s bioprocess prediction. To obtain this score, a mean genetic interaction profile was first computed across all *L*_2_-normalized genetic interaction profiles annotated to the biological process for which the inner product with the compound’s chemical-genetic interaction profile was 2 or greater. The importance score profile was then obtained by taking the Hadamard product (elementwise multiplication) between this mean genetic interaction profile and the compound’s chemical-genetic interaction profile.

**Overrepresentation analyses of gene mutants with strong chemical-genetic and/or genetic interactions**. After restricting the data to the top biological process prediction for each compound, gene mutants that possessed strong, negative chemical-genetic interaction scores (z-score < −5) were assessed for overrepresentation with respect to the number of times they did not contribute (importance score within ±0.1) to a compound’s top bioprocess prediction. Specifically, the number of times each strain occurred inside and outside the region described above (grey box in Figure 5) was compared to the number of times all strains occurred inside and outside the region using a hypergeometric test, using all strains with interaction z-scores < −5 as the background set. Details on the genes overrepresented in this region are given in Table S2.

### Experimental validation of compound-bioprocess predictions

**Phenotypic analysis of cell cycle progression**. To examine the effect of compounds on arresting cells in G2/M phase, we looked for differences in budding index and cell DNA content between compounds predicted to perturb the cell cycle versus negative control compounds. Seventeen compounds with high-confidence predictions to the bioprocess term “mitotic spindle assembly checkpoint” and strong negative chemical-genetic interactions with *PAT1* and *LSM6* (a common signature for compounds with this bioprocess prediction) were selected for validation. Additionally, ten bioactive (growth inhibition 50–80% compared to DMSO control) compounds with high confidence predictions (false discovery rate ≤ 25%) to bioprocess terms not related to cell cycle signaling and progression were selected as negative controls. Two compounds predicted to perturb “cell cycle phase” were also tested in these experiments. All compounds were tested at a concentration of 10 μg/mL, which was also the concentration used for chemical genomic screening [11].

To quantify budding index, logarithmically-growing *pdr1*Δ*pdr3*Δ*snq2*Δ cells were transferred to fresh galactose-containing medium (YPGal) containing compounds and incubated at 25 °C for 4 hours. The budding status of at least 200 cells was visually determined under the microscope. The percentage of the budded cells in no compound or compound-treated samples was counted.

For flow cytometry analysis, log phase *pdr1*Δ*pdr3*Δ*snq2*Δ cells were grown in YPGal media in the presence or absence of a compound for 4 hours; they were then fixed in 70% ethanol for 1 hour at 25 °C. Cells were collected by centrifugation, washed, and resuspended in buffer containing RNase A (0.25 mg/mL in 50 mM Tris, pH 7.5) for 1.5 hours. Cells were further incubated in 20 μl of 20 mg/ml proteinase K at 50 °C for 1 hour. Samples were then stained with propidium iodide, briefly sonicated, and measured using FACSCalibur ver 2.0 (Becton Dickinson, CA, USA).

The proportions of predicted active compounds and negative controls with positive phenotypic results were compared using the prop.test function in R to assess significance.

**Inhibition of tubulin polymerization**. *In vitro* tubulin polymerization assays using a fluorescent-based porcine tubulin polymerization assay (Cytoskeleton, BK011P) were performed following manufacturer specifications. Compounds were tested at a concentration of 10 μg/ml (with the exception of assay controls), which was identical to the concentration used for chemical genomic screening. All ten compounds predicted to perturb “tubulin complex assembly” with the minimum estimated false discovery rate (FDR < 1%) were selected for testing. Twelve compounds with predictions of false discovery rate ≤ 25% to any bioprocess except those related to chromosome segregation, kinetochore, spindle assembly, and microtubules were randomly selected as negative controls.

The degree of tubulin polymerization inhibition was summarized in a single *V*_max_ statistic for each compound treatment replicate. The *V*_max_ for each compound’s fluorescence time-course was calculated as the maximum change in fluorescence between consecutive time points, which were measured at 1-minute intervals. Three batches of experiments were performed in total (resulting in *N* ≥ 2 for each compound), and we normalized the *V*_max_ values in each batch by subtracting the difference between that batch’s mean DMSO (solvent control) *V*_max_ and the overall mean DMSO *V*_max_. To determine if the tubulin-predicted compounds inhibited polymerization to a significantly greater degree than the controls, we calculated the mean of the normalized *V*_max_ values for each compound and performed a one-sided Wilcoxon rank-sum to test for a difference in the ranks of these values between the two classes of compounds.

Chemical structure similarities between each pair of compounds selected for tubulin polymerization validation were obtained by first computing an all-shortest-paths fingerprint with path length 8 for each compound [44]. Similarities were computed on the fingerprints using the Braun-Blanquet similarity coefficient, which is defined as the size of the intersection divided by the size of the larger set. In a recent study, this combination of structure descriptor and similarity coefficient performed well when evaluated globally on our entire chemical-genetic interaction dataset [45].

## Acknowledgments

SWS would like to thank Hamid Safizadeh and Henry Neil Ward for their proofreading of the manuscript. This work was partially supported by the National Institutes of Health (https://www.nih.gov/) (R01HG005084, R01GM104975) and the National Science Foundation (https://www.nsf.gov/) (DBI 0953881). SWS is supported by an NSF Graduate Research Fellowship (00039202), an NIH Biotechnology training grant (T32GM008347), and a one-year fellowship from the University of Minnesota Bioinformatics and Computational Biology Graduate Program (https://r.umn.edu/academics-research/graduate-programs/bicb). SCL and JSP are supported by a RIKEN (http://www.riken.jp/en/) Foreign Postdoctoral Research Fellowship. SCL is supported by a RIKEN CSRS (http://www.csrs.riken.jp/en/) Research Topics for Cooperative Projects Award (201601100228), and a RIKEN FY2017 Incentive Research Projects Grant. YO is supported through Grants-in-Aid for Scientific Research (15H04402) from the Ministry of Education, Culture, Sports, Science and Technology, Japan (www.mext.go.jp/en/). CB and YO are supported by JSPS KAKENHI grant number 15H04483 (http://www.jsps.go.jp/english/). CB and YY are supported by a JSPS Grant-in-Aid for Scientific Research on Innovative Areas (17H06411). CLM and CB are fellows in the Canadian Institute for Advanced Research (CIFAR, https://www.cifar.ca/) Genetic Networks Program. Computing resources and data storage services were partially provided by the Minnesota Supercomputing Institute and the UMN Office of Information Technology, respectively. Software licensing services were provided by the UMN Office for Technology Commercialization. The funders had no role in study design, data collection and analysis, decision to publish, or preparation of the manuscript.

## Supporting figure legends

**Figure S1.** Performance comparison of CG-TARGET versus baseline enrichment approaches. Perturbed biological processes were predicted using both CG-TARGET and methods that calculated enrichment on the set of each compound’s *n* most similar genetic interaction profiles (“top n,” *n* ∈ {10, 20, 50, 100, 200, 300, 400, 600, 800}). (A) Biological process prediction false discovery rate estimates derived from resampled chemical-genetic interaction profiles, performed on compounds from the RIKEN dataset. (B) Precision-recall analysis of the ability to recapitulate gold-standard annotations within the set of top bioprocess predictions for ~4500 simulated compounds. Each simulated compound was designed to target one query gene in the genetic interaction network and thus inherited gold-standard biological process annotations from its target gene. (C) For each of 35 well-characterized compounds in the RIKEN dataset with literature-derived, gold-standard biological process annotations, we determined the rank of its gold-standard bioprocess within its list of predictions. The number of compounds for which a given rank (or better) was achieved is plotted. The grey ribbons represent the median, interquartile range (25^th^ to 75^th^ percentiles), and 95% confidence interval of 10,000 rank permutations.

**Figure S2.** Induced GO hierarchy of the 100 best-performing GO biological process terms, evaluated using simulated chemical-genetic interaction profiles. Each term was evaluated using precision-recall statistics (area under the precision-recall curve divided by the area under a curve produced by a random classifier) to analyze its ability to rank simulated chemical-genetic interaction profiles from which it was annotated as a gold-standard bioprocess. Green nodes represent the 100 best-performing GO biological process terms, yellow nodes represent terms for which predictions were made but did not rank among the top 100, and white nodes represent terms in the Biological Process ontology that were not selected for bioprocess prediction. Hovering the mouse over each node reveals its GO ID and name.

**Figure S3.** Induced GO hierarchy of the 100 worst-performing GO biological process terms, evaluated using simulated chemical-genetic interaction profiles. Same as Fig S2, but for the 100-worst performing GO biological process terms.

**Figure S4.** Schematic representation of CG-TARGET bioprocess prediction procedure. Further details on the presented procedures, including equations, are given in “Predicting the biological processes perturbed by compounds” in Materials and Methods.

## Supporting table legends

**Table S1.** Using protein complexes to refine CG-TARGET GO biological process mode-of-action predictions. Compounds, GO biological processes, and protein complexes are shown if the mode-of-action prediction to the protein complex was stronger than that to the associated GO biological process (comparison first based on p-value, then on z-score in the case of a tie). Protein complexes were limited to those of size 4 or greater whose gene annotations were a subset of those for the corresponding GO biological process term. The final column indicates compounds that did not achieve a false discovery rate of 25% or less for any GO biological process mode-of-action predictions but did for at least one protein complex prediction (with “HCS” denoting “high confidence set”).

**Table S2.** Overrepresentation analysis of mutant strains with strong negative chemical-genetic interactions and no contribution to top bioprocess predictions. Overrepresentation within the shaded region of Fig 5 was evaluated using a hypergeometric test to compare the occurrence of one strain versus all strains inside and outside of the region, with the background containing only strains that possessed strong (z-score < −5) negative chemical-genetic interactions. The compounds and top bioprocess predictions associated with each strain’s occurrences in the region are given, as well as the appropriate background list of strains and information on the gene deleted in each strain.

